# A defined growth medium for trace metal limitation studies in *Auxenochlorella sp.* (UTEX 250-A) during mixotrophic growth with glucose

**DOI:** 10.1101/2025.05.09.653207

**Authors:** Dimitrios J. Camacho, Jeffrey L. Moseley, Sabeeha S. Merchant

## Abstract

*Auxenochlorella sp.* (strain UTEX 250-A) is a fast-growing, oleaginous alga that is an emerging reference organism for basic research, with broad biotechnological applications. Advancing UTEX 250-A as a biotechnological workhorse requires a deeper understanding of its nutritional demands, particularly under different trophic conditions. However, little is known about the specific acclimations that allow UTEX 250-A to thrive when essential nutrients are scarce. Here, we describe the formulation of a defined growth medium that supports the cultivation of UTEX 250-A with controlled trace levels of the essential micronutrients iron, copper, and zinc. Special attention was given to ensure that the medium was compatible with inductively coupled plasma mass spectrometry (ICP-MS) analysis, enabling the detection and minimization of metal contamination in the nanomolar range. The medium was designed to provide sufficient nutrients to support the mixotrophic growth of UTEX 250-A from an initial density of 10^5^ cells/mL to a stationary density of 2.4×10^8^ cells/mL, with additional nutrients supplied to accommodate metabolic and trophic transitions during stationary phase, when photosynthesis is restored due to carbon limitation. This replete medium provides a foundation for robust nutrient limitation studies in *Auxenochlorella sp.* (UTEX 250-A).

## Introduction

The elemental composition of a cell is dynamically coordinated with its metabolic demands and the ratios of elements available in its environment. While many well characterized nutrient assimilation mechanisms are conserved across algae, and across organisms throughout the tree of life, less is known about organisms that have not been well-studied in the laboratory (Blaby-Haas & Merchant, 2012). *Auxenochlorella spp.* are trebouxiophyte algae with applications in waste-water remediation (Forghani et al., 2022; Gramegna et al., 2020; Heller et al., 2015; Talapatra & Ghosh, 2022; Zhou et al., 2012), fuel production (Franklin, Somanchi, et al., 2014; Patel et al., 2018), food production (Brooks et al., 2010; Canelli et al., 2020; Caporgno et al., 2020), and pigment production (Moseley et al., 2024; Xiao et al., 2018). It can switch between autotrophic and heterotrophic growth (Matsuka et al., 1966), and produces up to 58% of its dry weight as triacylglycerols during heterotrophy (Xiong et al., 2008). UTEX 250-A is of particular interest because of its robust growth, and it is an allodiploid hybrid closely related to *Auxenochlorella protothecoides* and *Auxenochlorella symbiontica* (Craig et al., 2025). Its compact genome and ease of genetic transformation via homologous recombination facilitates targeted reverse genetics approaches to investigate specific gene functions (Dueñas et al., 2025; Moseley et al., 2024). To disentangle the molecular circuits orchestrating the balance of UTEX 250-A’s elemental composition, we must first provide a controlled environment where all its nutritional requirements are met, except for the element that we selectively modulate. This isolation enhances the signal to noise ratio of specific cellular responses and prevents crosstalk from unintended deficiencies, facilitating more accurate insights from systems wide comparative multi-omics studies. In the case of metals like Fe, Cu, and Zn, which are required in trace amounts, precise modulation of their concentrations requires their complete exclusion from all other ingredients of the medium. This is because although they are essential, they are needed at very low concentrations in the cell (Merchant & Helmann, 2012). For instance, to achieve Cu deficiency in *Chlamydomonas reinhardtii* cells, all glassware and plasticware are rigorously washed with 6 M HCl and thoroughly rinsed with ultra-pure ICP-MS-grade water (Quinn & Merchant, 1998). The medium is prepared exclusively with ultra-pure reagents (>99.999% (w/w) purity) and cells are passaged three times in Cu deficient medium (<3 nM Cu) to deplete their intracellular Cu stores (Kropat et al., 2015).

The existing UTEX 250-A growth medium, ApM1, was optimized for the maximum production of lipids during heterotrophic growth (Franklin, Brubaker, et al., 2014). However, it is not compatible with typical inductively coupled plasma mass spectrometry (ICP-MS) instrumentation due to high levels of sodium, potassium, and phosphorous (**Figure 1**). When macronutrients are provided in such quantities, the potential contribution of metal impurities from a given component increases. The first ionization potentials (IE_1_), or ionization energies required to form singly positively charged ions, of sodium and potassium are low (5.14 and 4.34 electron volts (eV), respectively) (Ciocca et al., 1992; Kramida & Ralchenko, 1999; Lorenzen et al., 1981). In contrast, the first ionization potentials of the trace metals in which we are interested are relatively high (**Table 1)**. The ionization potential of an element is inversely correlated with its ionization efficiency. Due to the high concentrations of Na (26 mM) and K (48 mM) in the ApM1 medium (**Figure 1**), a substantial fraction of the inductively coupled plasma’s ionization energy would be directed towards ionizing Na and K, leaving less energy available for the ionization of other target elements with higher ionization potentials, a phenomenon known as ion suppression (Houk et al., 1980; Kalnicky et al., 1977; Olivares & Houk, 1986). A 40-fold dilution of the ApM1 medium is needed to bring the K concentration into the calibration range of most ICP-MS instruments, just below the upper threshold of 50 ppm or 1.3 mM K. This dilution may also decrease the concentration of trace elements below quantitation limits during growth. For example, replete ApM1 medium contains 441 nM of Cu; diluted 40-fold would yield 11 nM Cu. The molar concentration of K in ApM1 is more than 100,000-fold higher than that of Cu. The typical lower limit of quantitation for Cu using our instrument is around 1.6–3.2 nM Cu, but only in ideal conditions when Cu ionization is not suppressed.

**Figure 1.**
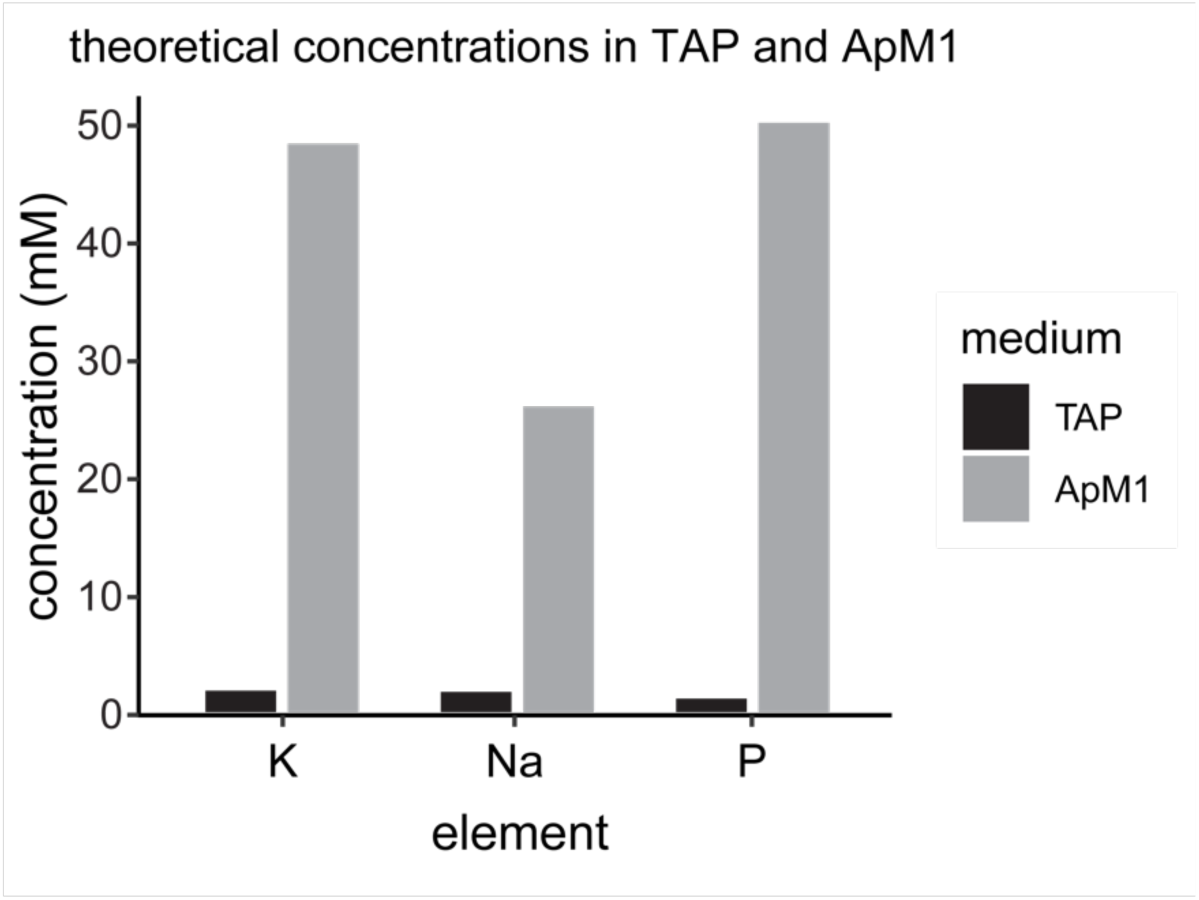
Theoretical concentration of Na, K, and P in TAP and ApM1 media.

**Table 1.** First ionization potential of select elements (Kramida & Ralchenko, 1999) element first ionization.

| element | first ionization<br>potential (eV) |
| --- | --- |
| C | 11.26 |
| Na | 5.14 |
| P | 10.49 |
| S | 10.36 |
| K | 4.34 |
| Fe | 7.90 |
| Cu | 7.73 |
| Zn | 9.39 |

Other strategies for accurately and precisely measuring media with high total dissolved solids content by ICP-MS do exist, but they are costly and time consuming. One approach is through standard addition and involves adding the sample to a series of standards with increasing concentrations that are within the calibration range to generate a separate standard curve (Camacho, Perrino, et al., 2024a). However, this method significantly extends instrument run time, as it must be repeated for each standard associated with a given sample. A three-point calibration for each sample would increase the analysis time three-fold. Argon (Ar) is the most expensive reagent for the operation of ICP-MS/OES instruments. The analysis of one sample takes roughly 7 min, during which 105 L of Ar gas is consumed. In addition, each sample requires dilution into a new tube, which increases waste, reagents, consumables, and introduces risks of contamination and user error. A more effective approach for quantifying trace elements in samples with a high total dissolved solids content by ICP-MS is the use of an ultra-high matrix introduction system provided by Agilent (Proper et al., 2014). Nevertheless, routinely running samples with high total dissolved solids content physically taxes the components of the ICP-MS instrument. For example, the deposition of salts on the sample and skimmer cones may cause the sensitivity of the instrument to drift, even over the course of a single analysis session. The frequency of cleaning and instrument downtime will increase. Sample introduction components, such as tubing, nebulizers, and spray chambers can accumulate residue over time, potentially leading to the sporadic release of contaminants and confounding results. The introduction of more chemical species also increases the chances of polyatomic ion interferences, which may present a multitude of complications in data analysis. The data presented here are not exempt from these challenges. Both ion suppression and ion enhancement occurred in many samples, and we employed strategies to minimize matrix effects and normalize the data. For instance, samples are co-injected into the plasma with the internal standards yttrium (^89^Y), scandium (^45^Sc), and rhenium (^187^Re). The recoveries of these internal standards often diverge when high matrix samples are analyzed. The target analyte should be normalized to the appropriate internal standard with the most comparable first ionization potential and atomic mass, as these factors influence how different ions behave during plasma ionization and ion transport (Finley-Jones et al., 2008; Wilschefski & Baxter, 2019). However, the behavior of target analyte ions is not always predictable in complex matrices, and internal standard normalization alone may not correct the full extent of matrix effects. The purpose of this optimization was to provide UTEX 250-A with a nutrient replete medium, while ensuring accurate quantification of its components. Here, we discuss the limitations of initial approaches, the obstacles encountered, and the solutions we employed to optimize the medium.

## Methods

### Strains and culture conditions

The wild-type *Auxenochlorella sp.* strain UTEX 250-A was derived from a single colony of UTEX 250, which was originally acquired from the University of Texas-Austin Culture Collection of Algae (UTEX). UTEX 250-A cells were grown in various media types, but all supplemented with 2% (w/v) glucose. All UTEX 250-A cultures were grown in 250 mL flasks containing 100 mL of liquid medium. The flasks were agitated at 200 rpm and held at 28 °C in a Multitron Pro incubator (Infors HT, Bottmingen, Switzerland) and illuminated continuously with 100 µ mol photons m^-2^ s^-1^ using warm white (3435 K) LED lights. For each experiment, three biological replicates defined as individual culture flasks, inoculated from starter culture, were cultivated and measured simultaneously. All experimental cultures were inoculated from a mid-log phase culture containing the HPv1 medium described in **Table 2**. Growth was assessed by measuring the cell density of each culture using a hemocytometer. A detailed protocol for counting cells with a hemocytometer is described in (Camacho & Merchant, 2024). Flasks in figures 2, 12, and 16 were photographed following a protocol described in (Camacho, Glaesener, et al., 2024).

**Figure 2.**
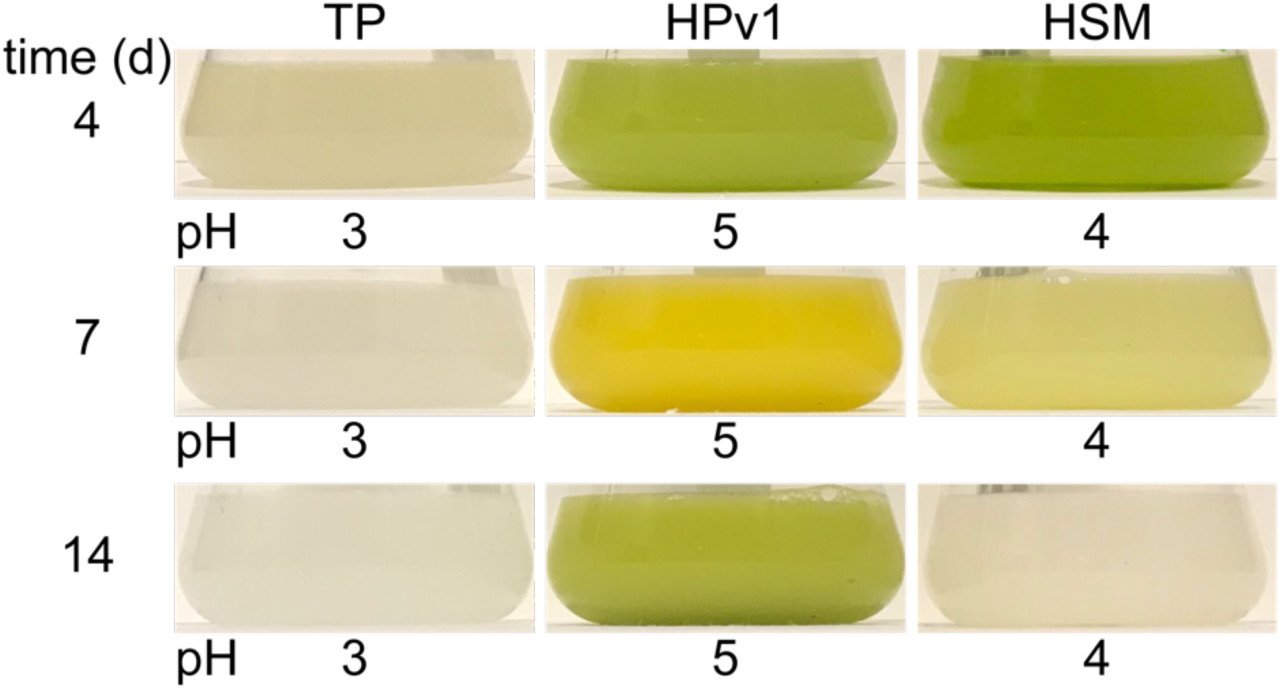
Growth of UTEX 250-A supplemented with 2% glucose in TRIS-phosphate (TP), HEPES-phosphate version 1 (HPv1) and high salt minimal (HSM) media. Cl^−^ was used as a counter-ion for TRIS, while Na was used as a counter ion for HEPES.

**Table 2.** Composition of the HEPES-phosphate medium versions that were tested throughout growth. Differences between the previous version are highlighted in yellow.

|  | species | TAP | HPv1 | HPv4 | HPv6 | HPv7 |
| --- | --- | --- | --- | --- | --- | --- |
| (mM) | NH<sub>4</sub>Cl | 7.5 | 7.5 | 7.5 | 7.5 | 15 |
|  | CaCl<sub>2</sub> | 0.3 | 0.3 | 0.3 | 0.3 | 0.7 |
|  | MgSO<sub>4</sub> | 0.4 | 0.4 | 0.4 | 1.2 | 2.4 |
|  | HEPES | 0 | 20 | 20 | 10 | 10 |
|  | K<sub>2</sub>HPO<sub>4</sub> | 0.7 | 0.7 | 0.7 | 6.8 | 6.8 |
|  | KH<sub>2</sub>PO<sub>4</sub> | 0.5 | 0.5 | 0.5 | 4.5 | 4.5 |
| (μM) | Zn | 2.5 | 2.5 | 20 | 20 | 20 |
|  | Mn | 6 | 6 | 18 | 18 | 18 |
|  | Fe | 20 | 20 | 100 | 100 | 100 |
|  | Cu | 2 | 2 | 15 | 15 | 15 |
| (nM) | Mo | 28.5 | 28.5 | 28.5 | 28.5 | 28.5 |
|  | Se | 1 | 1 | 1 | 1 | 1 |

### Transmission Electron Microscopy (TEM)

Transmission Electron Microscopy was performed using protocols developed by Dr. Radhika Mehta, Dr. Danielle Jorgens, and Reena Zalpuri. Briefly, 9 mL of a logarithmically growing UTEX 250-A (3 × 10^7^ cells/mL) culture in HPv11 medium, described in (Camacho, Perrino, et al., 2024c), was centrifuged at 17,000 ×*g* in a Beckman Coulter Avanti JXN-26 centrifuge equipped with 15 mL adapters for the fixed angle JA-14.50 centrifuge rotor. 6.44 mL of the supernatant was transferred to a new 15 mL falcon tube and 0.56% of glutaraldehyde was added to the supernatant. The mixture was wrapped in foil and inverted to mix. 7 mL of the logarithmically growing UTEX 250-A (3 × 10^7^ cells/mL) culture was added to the supernatant and glutaraldehyde mixture and shaken in the dark for 1 h at room temperature using a Thermolyne M26125 shaker. The sample was brought to the UC Berkeley Electron Microscopy Lab (EML) for further processing with help from Reena Zalpuri. The sample was quickly centrifuged for 1 min using an Eppendorf 5412 constant speed centrifuge. All subsequent centrifugation steps were carried out using this centrifuge at variable durations to achieve tight cell pellets. The supernatant was removed, and the cell pellet was resuspended in 1 mL of 1% OsO_4_ in 1×PBS, then shaken for 1 h in the dark at room temperature. The addition of 1.6% potassium ferricyanide was omitted. The sample was centrifuged to remove the supernatant, and the resulting pellet was washed five times by resuspension in 500 µL of 1× PBS and centrifugation. After the final wash, the pellet was resuspended in 1 mL of 1× PBS and stored overnight at 4 °C.

For dehydration, an ethanol gradient was used in place of acetone. The sample was centrifuged again to discard the supernatant, and the cell pellet was resuspended in 1 mL of 35% of ethanol. The suspension was gently mixed by inversion on a rotating mixer for 10 min. The sample was centrifuged to pellet the cells, which were subsequently resuspended in 50% ethanol using a toothpick. The process of resuspension in ethanol, inversion for 10 min, and centrifugation was repeated sequentially using 75%, 90%, and two rounds of 100% ethanol. The sample was centrifuged, and the supernatant was discarded.

The cells were infiltrated through three gradients of acetone:Epon resin (1:2, 1:1, 2:1). The Epon resin was made with 23.5 g Eponate 12 resin, 12.5 g dodecenyl succinic anhydride, 14 g nadic methyl anhydride, 0.750 mL benzyldimethylamine as the accelerator. The pure resin with accelerator was incubated in a vacuum chamber for 15 min to remove air bubbles. The dehydrated sample was gently resuspended in 1 mL of 1:2 resin:acetone mixture using a toothpick and mixed on a rotating mixer for 1 h. The sample was centrifuged until a dark cell pellet was visible and the supernatant containing resin was discarded. The sample was gently resuspended in 1 mL of 1:1 resin:acetone using a toothpick and mixed by inversion for 1 h on a rotating mixer. The sample was centrifuged until a pellet was visible and the supernatant was discarded. The sample was resuspended in 1 mL of 2:1 resin:acetone gently using a toothpick and was inverted to mix for 1 h. Next, the sample was centrifuged, and the supernatant was removed. The pellet was resuspended in 1 mL of pure resin (with accelerator) and inverted for 30 min before centrifugation and removal of the supernatant. This step was repeated. A small amount of resin was added to the pellet and cells were transferred into round bottom, flat-top capsules. The cells were concentrated to the bottom of the tube by centrifugation at 12,000 ×*g* until a loose pellet became visible. The resin was cured in a 65 °C oven for 48 h.

Reena Zalpuri performed the microtome sectioning of the cells embedded in resin (70–100 nm thin sections) and placed the sections onto molybdenum TEM grids. Sections were stained with lead citrate and uranyl acetate. Briefly, uranyl acetate was centrifuged for 10 min and added to a gid holder containing Mo TEM grids. Grids were incubated for 7 min, removed from the uranyl acetate mixture, and sequentially dipped into four beakers of Milli-Q H_2_O. Pb citrate was filtered through a 0.22 µm pore size syringe filter and added to the grid holder containing the TEM grids. Grids were incubated in Pb citrate for 5 min followed by four sequential washes in Milli-Q H_2_O. Dr. Jorgens and Reena Zalpuri initiated the Tecnai 12 Transmission electron microscope (120 kV with Rio16 CMOS camera, Gatan Inc, CA, USA) and loaded the grids on the specimen holder and into the column. Dr. Jorgens, Dr. Mehta, and Reena Zalpuri provided training and supervision during TEM image acquisition.

### Elemental analysis of the spent medium

Samples for elemental analysis were prepared following a protocol outlined in (Camacho, Perrino, et al., 2024a). Briefly, approximately 10^8^ cells were collected by centrifugation (17,000 ×*g*, 2 min) using a Beckman Coulter Avanti JXN-26 centrifuge with 15 mL adapters in a fixed angle JA-14.50 centrifuge rotor. 0.5 mL of the supernatants containing spent media were transferred to new tubes. 1.5 mL of water was added to each tube along with 5 mL of 2.8% HNO_3_. The spent medium was measured by inductively coupled plasma mass spectrometry (ICP-MS/MS) following a protocol outlined in (Camacho, Perrino, et al., 2024b). Briefly, elements were measured from samples using either H_2_, He, or O_2_ tune modes. Each element was normalized to one of three elements in an internal standard containing Sc (Agilent 5190-8517), Y (5190-8555), and Re (5190-8507). The instrument was calibrated using the environmental standard (Agilent 5183-4688), P calibration standard (Inorganic Ventures CGP1), and S calibration standard (Inorganic Ventures CGS1).

### Total organic carbon (TOC) measurements of UTEX 250-A cells

Approximately 10^8^ UTEX 250-A cells cultivated in HPv1 (**Table 2**) with 2% (w/v) glucose were collected every 12 h throughout growth to stationary phase by centrifugation (17,000 ×*g*, 2 min). The supernatants were discarded, and the pellet was washed once with 1 mL of 1 mM Na_2_EDTA and once with 1 mL of Milli-Q water. The cell pellets were digested in 286 µL of 70% HNO_3_ and 10 mL of Milli-Q H_2_O was added for a final matrix of 2% HNO_3_. 500 µL of digested and diluted cell material was added to a TOC measurement vial containing 14.5 mL of Milli-Q H_2_O. The sample was measured by a Shimadzu Total Carbon Analyzer. Non-purgeable organic carbon (NPOC) was used as a proxy for biomass and was normalized to mL of culture sampled (µg NPOC/mL of culture) and cell number (pg NPOC/cell number).

## Results

The ideal medium would contain only those nutrients necessary to sustain the rapid mixotrophic growth of *Auxenochlorella* UTEX 250-A (cultivated with 2% (w/v) glucose under 100 µmol photons m^−2^ s^−1^ illumination) and enough of those nutrients to allow cells to reach stationary phase. It should be chemically defined, containing only ultra-high purity trace metal grade chemicals, tested by the manufacturer, and accompanied by certificates of analysis for each specific batch, with minimal batch to batch variation. The components of the medium should be easy to obtain, economical, simple to prepare, and should have a long shelf-life. The components of the medium should not cross-react, form highly stable complexes with metals, nor should they cause other nutrients in the medium to precipitate. The medium should provide sufficient buffering capacity, with a buffering species p*K*_a_ between 6 and 8. Ideally none of the components would be toxic if ingested or inhaled. Lastly, and perhaps most importantly, it must be compatible with a wide variety of experimental protocols and analytical instruments.

As a starting point, we chose to base the medium composition on the already established tris-acetate-phosphate (TAP) medium, used to grow *Chlamydomonas reinhardtii* (Gorman & Levine, 1965; Hui et al., 2023), supplemented with the “Special K” trace element solution (Kropat et al., 2011). TAP medium already satisfied most of our requirements and our ICP-MS instrument was previously calibrated to precisely measure its elemental composition with minimal polyatomic ion interferences. Importantly, UTEX 250-A grows in TAP medium.

### Optimization of the buffer system for mixotrophic growth of UTEX 250-A in 2% (w/v) glucose

Although UTEX 250-A metabolizes acetate, glucose supports more a rapid growth and is commonly used as a carbon source in commercial scale fermentations to generate valuable natural products (Brooks et al., 2010). Despite the substantially large reduced carbon concentration (666 mM) from 2% (w/v) (111 mM) glucose supplementation, the effect of ion suppression on target analytes is less of a concern due to carbon’s high first ionization potential (11.26 eV) (Haris & Kramida, 2017; Kramida & Ralchenko, 1999). However, this does not entirely eliminate its effect on plasma dynamics. The accelerated growth facilitated by glucose supplementation comes at the expense of rapid nutrient depletion and acidification of the culture medium (**Figure 2**). As UTEX 250-A cells metabolize ammonium chloride (NH_4_Cl), they release protons and acidify the medium (Wang & Curtis, 2016). In other systems, a dual nitrogen source in the form of ammonium nitrate is used to achieve pH control (Imsande, 1986; Scherholz & Curtis, 2013). However, UTEX 250-A is incapable of utilizing nitrate (NO_3_^−^) as a nitrogen source; thus, the usage of NH_4_NO_3_ would fail to balance the pH. In standard TAP medium, buffering is provided by 20 mM TRIS (p*K*_a_ of 8.1) titrated with 17 mM acetic acid to pH 7.0–7.2, in combination with a balanced phosphate buffer system comprising of monobasic (0.45 mM KH_2_PO_4_, relevant p*K*_a_ 2.15 and 7.21) and dibasic (0.68 mM K_2_HPO_4_, relevant p*K*_a_ 7.21 and 12.38) potassium phosphate. TRIS is particularly suited for use with acetate, as the assimilation of acetic acid leads to medium alkalization, reaching up to pH 8.5 at stationary phase in *Chlamydomonas* (Hui et al., 2022). When acetate is omitted from TAP medium for photoautotrophic growth (in TP medium), TRIS base is protonated using HCl. When 17 mM acetate was substituted for 2% (w/v) glucose in UTEX 250-A cultures, the pH of the medium dropped from pH 7 to pH 3 after four days of growth to stationary phase cell densities (**Figure 2**). In the first round of media optimizations, we chose to substitute 20 mM of TRIS-acetic acid for 20 mM of HEPES titrated with NaOH to pH 7 and named this medium HEPES-phosphate version 1 (HPv1). Na-HEPES was chosen as a replacement buffer because it has been generally regarded as a non to very weak metal complexing agent (Ferreira et al., 2015; Good et al., 1966). Its p*K*_a2_ of 7.55 enables it to effectively maintain a pH within a range of 6.8 to 8.2. Replacing TRIS with Na-HEPES in glucose supplemented cultures allowed pH stabilization at pH 5 after cells reached a stationary phase density of ∼2.4×10^8^ cells/mL on day 4 (**Figure 2**). In parallel, we cultured UTEX 250-A in a high salt medium (HSM) (Sueoka, 1960), a standard minimal medium for *Chlamydomonas* growth that utilizes an increased potassium phosphate concentration as its sole buffering system (8.25 mM K_2_HPO_4_ and 5.25 mM KH_2_PO_4_). When HSM grown UTEX 250-A cultures were supplemented with 2% glucose, we also observed acidification of the medium after 4 d, though to a lesser extent than TP, reaching pH 4 instead of pH 3. Both TP and HSM cultures appeared lighter in color than the HPv1 culture after 7 d (**Figure 2**). The HSM culture eventually turned white after 14 d.

### Determining the elemental quota of UTEX 250-A on a per cell basis presents technical challenges

With the pH stabilized, we then asked what the elemental demands of UTEX 250-A might be when grown mixotrophically with 2% (w/v) glucose and in continuous light (100 µmol photons m^−2^ s^−1^) with 200 rpm agitation, at 28 °C. To answer this question, we measured the concentrations of Na, Mg, P, S, K, Ca, Mn, Fe, Cu, Zn, Se, and Mo in logarithmically growing cells and attempted to calculate the concentrations of each element required in the medium to sustain the growth of a culture to the stationary density of 2.4×10^8^ cells/mL. Collecting nutrient replete cells during the early logarithmic growth phase was essential to assess the intracellular elemental content prior to onset of potential nutrient limitation.

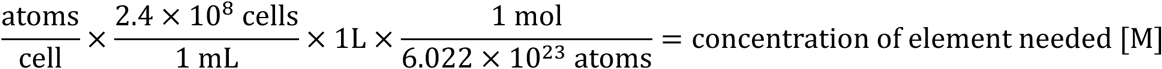

**Equation 1.** Calculation of element concentrations required to sustain a UTEX 250-A culture to stationary phase cell densities.

Determining the elemental demands of a UTEX 250-A cell was challenging due to the extensive variation in cell sizes within these unsynchronized batch cultures (**Figure 3a**). During logarithmic growth, cells increased in size and divided into multiple smaller daughter cells by multiple fission, resulting in a heterogenous population (**Figure 3b**, **3c**). To account for this variability, total organic carbon (TOC) was measured as a proxy for biomass, capturing fluctuations in biomass normalized to cell number throughout logarithmic growth phase (12–72 h) (**Figure 4**). The cell densities of cultures remained relatively unchanged from 12 h to 24 h (**Figure 4a**). However, the biomass of the cells increased sharply (**Figure 4c**). As a result, calculating the elemental demands of UTEX 250-A based on data from cells collected at 12 h versus 24 h would yield different results if calculated on a per cell basis.

**Figure 3.**
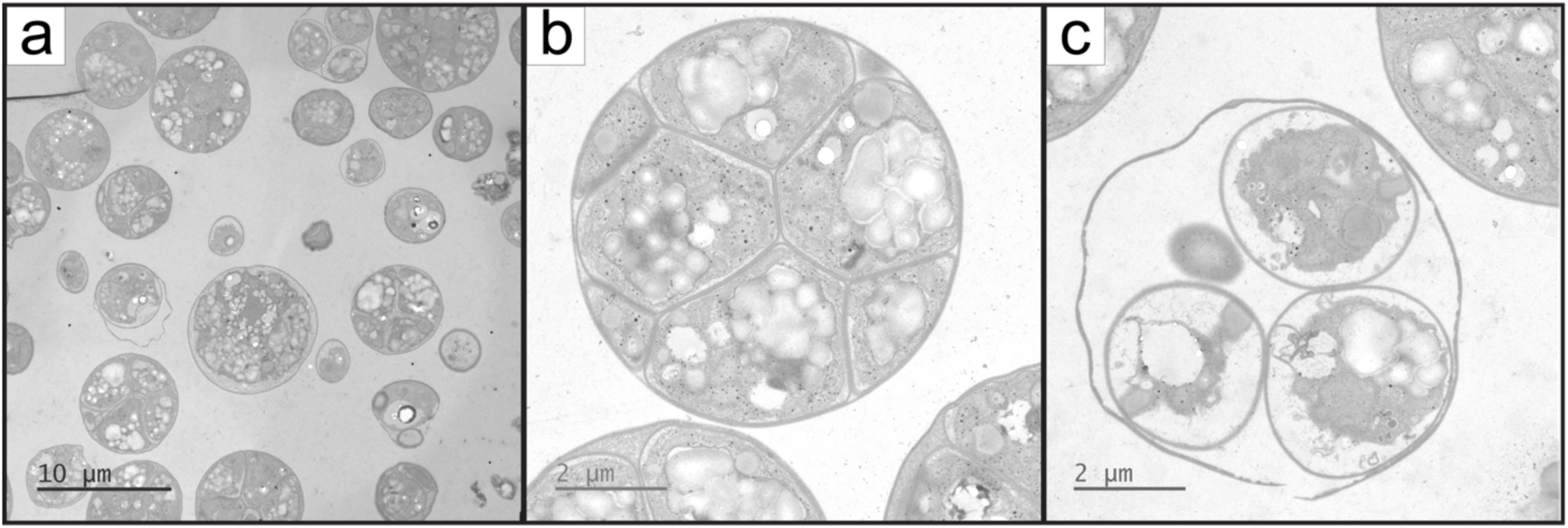
TEM images of UTEX 250-A cells undergoing multiple fission. Cells were grown in HPv1 supplemented with 2% (w/v) glucose under light conditions (100 µmol photons m^−2^ s^−1^) and sampled at the logarithmic growth phase (3×10^7^ cells/mL) for TEM analysis. **a)** Cells of varying sizes are shown. **b)** A single cell prior to multiple fission. **c)** The release of four daughter cells, with only a small portion of the fourth cell visible, likely due to its position outside the plane of sectioning during microtome slicing.

**Figure 4.**
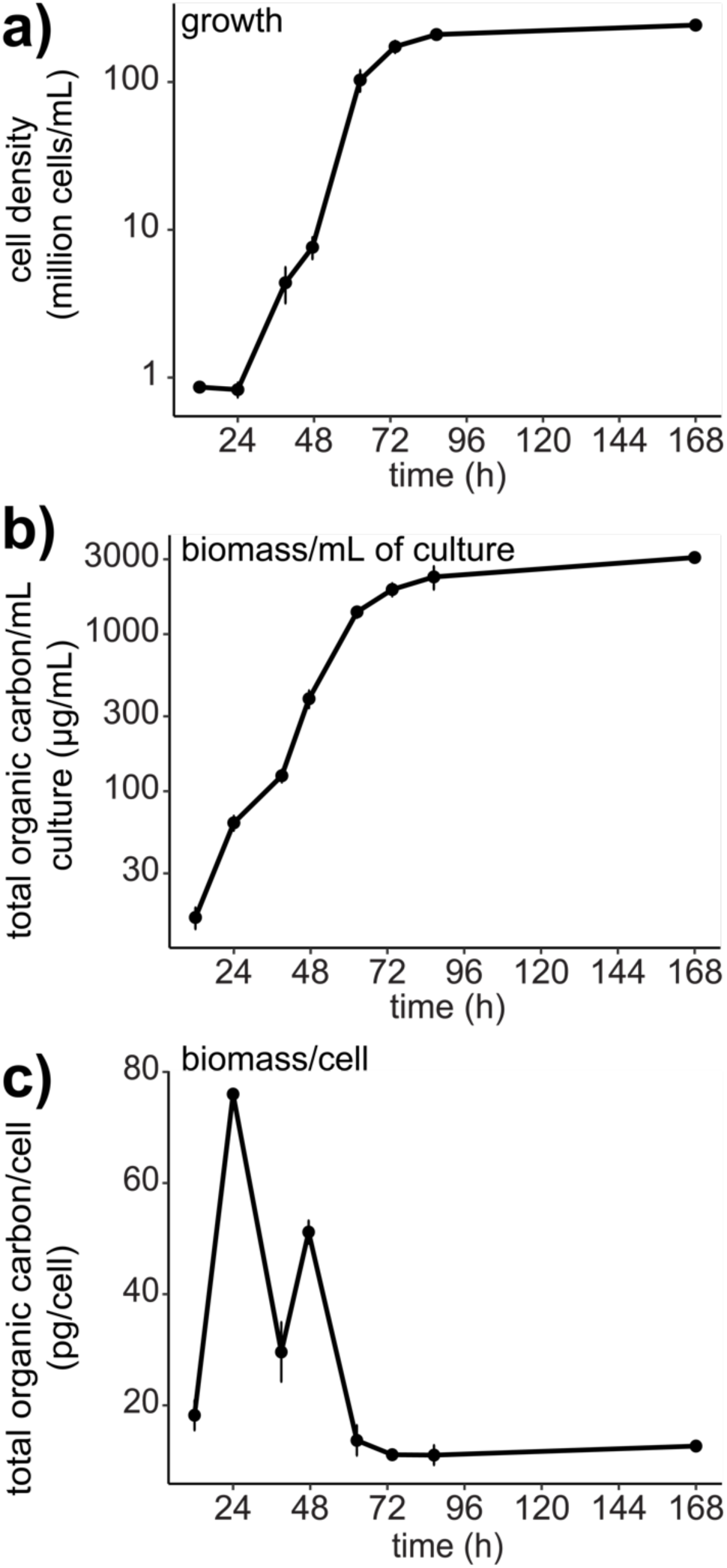
Variation in biomass of logarithmically growing UTEX 250-A cells. UTEX250-A cells were cultivated in 250 mL Erlenmeyer flasks containing HPv1 medium supplemented with 2% (w/v) glucose and placed in illuminated incubators (100 µmol photons m^−2^ s^−1^) at 28°C and agitated at 200 RPM. Error bars represent the standard deviation between three replicate culture flasks. **a)** UTEX 250-A growth as determined by a hemocytometer. **b)** Biomass (non-purgeable organic carbon (NPOC)) normalized to mL of culture sampled. Y-axis is log scaled. **c)** biomass normalized to cell number. The error bars do not indicate the standard deviations of cell-to-cell variation in biomass but rather the variation between individual replicate flask measurements.

The cells eventually become homogenous in biomass at around 10–13 pg/cell when division slows, and the culture enters stationary phase (**Figure 4c**). However, an additional layer of complexity arises from the ability of some algal cells to accumulate and sequester certain elements beyond their immediate growth requirements, particularly in response to stress or metal deficiencies, which may occur at late logarithmic to stationary phases. For example, Zn deficient *Chlamydomonas* cells have been shown to accumulate Fe, Mn, and Cu levels exceeding those in replete cells (Hong-Hermesdorf et al., 2014; Malasarn et al., 2013).

A key challenge in this analysis is the need for a highly accurate and absolute quantification of elemental content in cell samples, which requires 100% recovery of all cells within a sample tube. This was difficult, given the presence of smaller UTEX 250-A cells (>2 µm in diameter (**Figure 3**) relative to *Chlamydomonas reinhardtii* cells (10 µm). Smaller cells typically have less mass and a higher surface area to volume ratio; thus, they require increased centrifugal forces applied over longer durations for recovery. Additionally, all reagents and plastics need to be absolutely free of contaminants. To enhance cell recovery, we used special metal free tubes (Cat. No. 6295, Globe Scientific) with higher maximum centrifugal force ratings (17,000 ×*g*) in a Beckman Coulter Avanti JXN-26 centrifuge equipped with 15 mL adapters for the fixed angle JA-14.50 centrifuge rotor. These modifications allowed us to increase the centrifugal forces applied by 13,780 ×*g* compared to previous methods used for *Chlamydomonas* (3,220 ×*g*).

### Analysis of the HPv1 spent medium during growth provides insights into the timing of nutrient assimilation in UTEX 250-A

To circumvent the challenges of the approach described in **Equation 1**, we chose to analyze the elemental content of the spent medium every 12 h throughout growth to stationary phase and again at 7 days post-inoculation, to ensure that the cells remained replete with sufficient nutrients (except carbon and nitrogen). Three biological replicates were cultivated in separate flasks and their spent media was measured by ICP-MS. This provided a more direct approach for determining which nutrients were assimilated during growth and the timescales over which they were depleted in the medium. This analysis revealed that the rate of P assimilation plateaued around 60 h, preceding the transition to stationary phase 24 h later at 84 h, suggesting that P may have been a limiting nutrient (**Figure 5b**). Cells did not assimilate P to below 10–20 µM P. Like P, the rate of Mn assimilation diminished after 60 h. A similar observation was found for Mg, albeit later at 84 h. Zn levels in the spent medium declined beyond the lower quantitation limit (400 nM) of this analysis and could not be accurately quantified after 84 h, suggesting that the cells had insufficient Zn. The levels of all other elements measured remained above the quantitation limits, even after 7 days. However, it was unclear whether UTEX 250-A possessed the capacity to assimilate nutrients below the concentrations remaining in the medium at the point where the rate of assimilation plateaued. It could be interpreted as the cessation of assimilation being due to an inherent limitation in high-affinity assimilation, or simply because the cells no longer required additional nutrients. Specific nutrient limitation experiments would be necessary to determine the lower threshold of high affinity uptake pathways, but before this, the possibility of co-limitations must first be ruled out.

**Figure 5.**
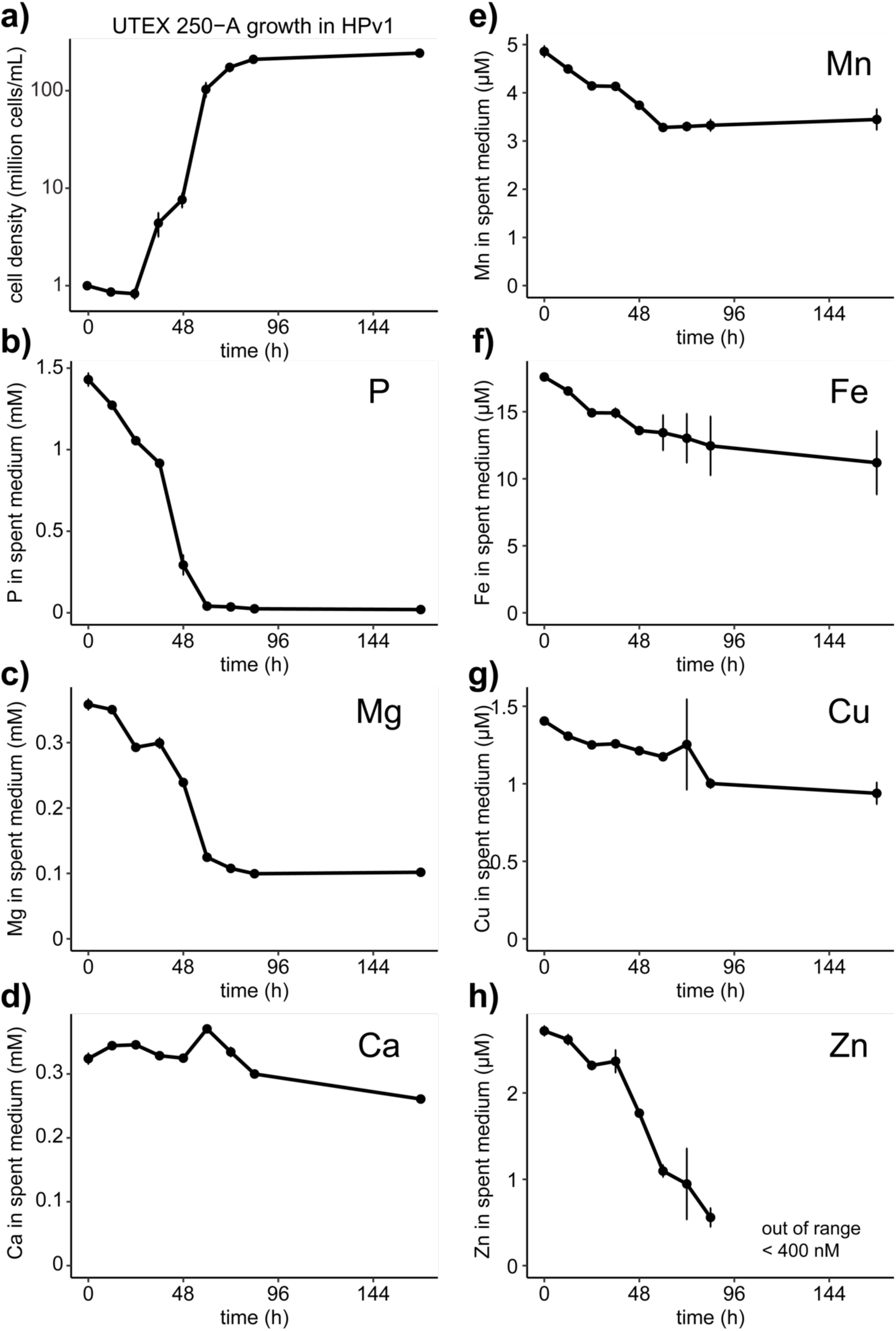
Nutrient consumption of *Auxenochlorella sp.* (UTEX 250-A) from the HPv1 medium supplemented with 2% (w/v) glucose. Error bars represent the standard deviation between three replicate flasks cultivated simultaneously from the same starter culture. **a)** Growth of UTEX 250-A over a period of seven days in illuminated incubators (100 µmol photons m^−2^ s^−1^). Cell density was measured using hemocytometer and the spent medium was sampled for elemental analysis every 12 hours until the cells reached stationary phase (2–2.5×10^8^ cells/mL).

To address the possible co-limitation of Mg and P in HPv1 we attempted to increase MgSO_4_ and P content by 10-fold in HPv2. We wondered if the 10-fold increase in the K_2_HPO_4_ and KH_2_PO_4_ buffer system would be sufficient to maintain the pH of the medium. The replacement of TRIS-acetate (C_4_H_11_NO_3_-C_2_H_3_O_2_) with Na-HEPES (Na-C_8_H_17_N_2_O_4_S) increased the S and Na concentrations of HPv1. We worried about the effects of additional S and Na on the ICP-MS instrument and chose to lower the Na-HEPES concentration to half the HPv1 concentration in HPv2. The S from MgSO_4_ is bioavailable whereas the S from Na-HEPES is not metabolized. We also formulated HPv3 simultaneously such that the only difference from HPv2 was the complete omission of Na-HEPES. Unfortunately, precipitates formed in HPv2 and HPv3 after autoclave sterilization. We found that adding NH_4_Cl, CaCl_2_, and MgSO_4_ prior to autoclavation caused the precipitation (**Figure 6**). We attempted to add the filter sterilized mixture of NH_4_Cl, CaCl_2_, and MgSO_4_ after autoclave sterilization and although no precipitates immediately formed, they became visible after three days at room temperature and would not dissolve back into solution with agitation. We were not able to use HPv2 and HPv3.

**Figure 6.**
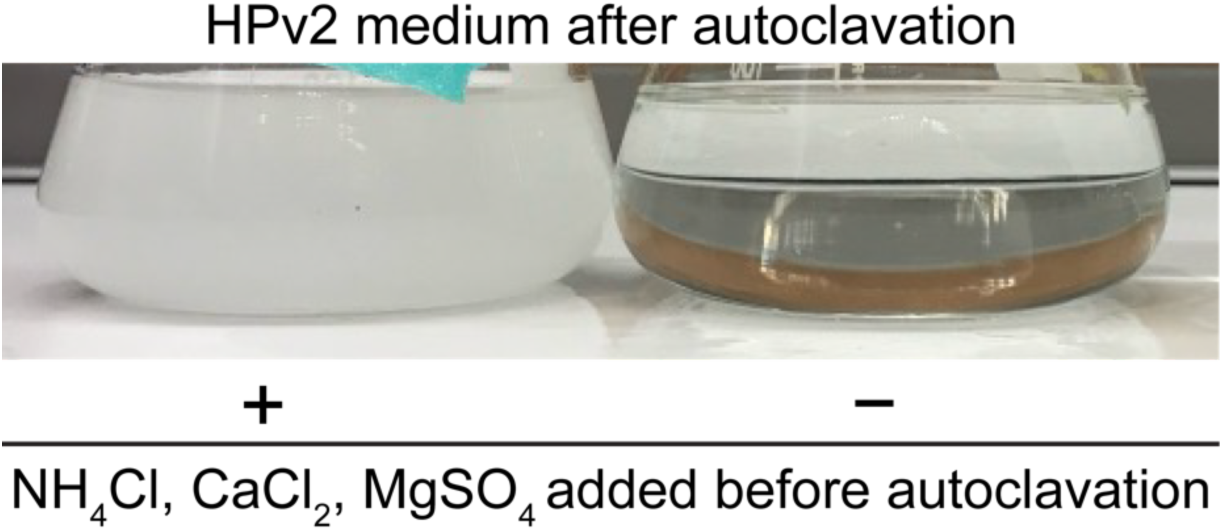
Precipitation of HPv2 medium after autoclave sterilization. A modified Beijerincks solution was added to one flask (left) before autoclave sterilization.

To raise MgSO_4_ and P levels without causing precipitation after autoclavation, we formulated HPv5, containing 1.2 mM of MgSO_4_ as opposed to 4 mM in HPv2 and HPv3. The Na-HEPES buffer was omitted, and the P buffer system was increased from 1.13 mM total P in HPv1 and HPv4 to 6.5 mM total P in HPv5. HPv5 also precipitated after autoclavation and was not investigated further (**Figure 7**).

**Figure 7.**
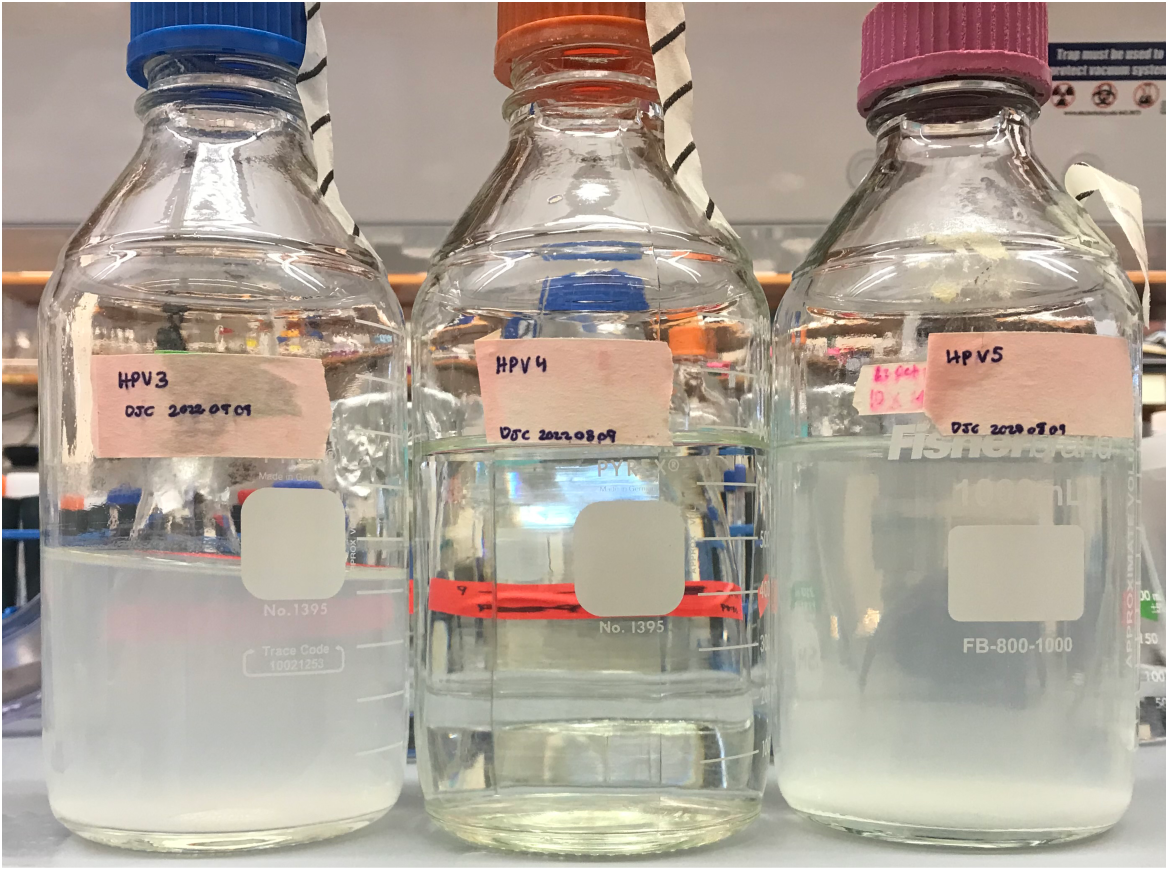
Precipitation of HPv3 (left) and HPv5 (right) media after autoclave sterilization. The HPv4 medium (center) did not form precipitates after autoclave sterilization.

Like HPv5, HPv6 featured an increased MgSO_4_ concentration (1.2 mM). We reduced the Na-HEPES concentration from 20 mM to 10 mM and increased the potassium phosphate buffer system to a total of 11.3 mM P. The stock salt solution, a modified Beijerink’s solution (Sueoka, 1960), contained NH_4_Cl, CaCl_2_, and MgSO_4_. The order in which these salts are combined also influences their solubility in the stock solution. A 100× concentrated stock salt solution led to precipitation, so we opted for a 40× stock salt solution instead, adding the filter sterilized stock after autoclaving the medium. This strategy enabled HPv6 and all subsequent versions to remain free of precipitates when stored at room temperature after autoclave sterilization. We also considered whether the cells were nitrogen limited, so we doubled the NH_4_Cl concentration in HPv7 to match the N content of the ApM1 (15 mM). HPv7 also contained 2-fold more CaCl_2_ and MgSO_4_ than HPv6. We knew that additional NH_4_Cl would acidify the medium but wanted to test the buffering capacity of the combined Na-HEPES and phosphate buffer system.

We wondered if the accumulation of P, Ca, K, and Mg in cells were a symptom of trace metal insufficiencies, so we formulated HPv4, which is identical to HPv1 except for increased levels of the trace elements Mn, Fe, Cu, and Zn (**Table 2**). If the accumulation of P, Ca, K, and Mg in cells were a symptom of Mn, Fe, Cu, or Zn deficiency, then we would expect P, Ca, K, and Mg to remain at higher concentrations in the Hpv4 spent medium after growth. The trace element concentrations chosen were initially based on calculations using **Equation 1** and remained unchanged from HPv4–HPv7.

We then cultured UTEX 250-A in HPv1, HPv4, HPv6, and HPv7 media supplemented with 2% (w/v) glucose and measured growth, pH of the spent medium, and the elemental composition of the spent medium every 12 h for 84 h. We also obtained measurements after 7 days (168 h) to determine if the cellular elemental composition was stable and not deficient in any element except for C and N (which we did not measure) at stationary phase. Three replicate flasks were cultivated for each medium, and we observed similar growth rates across the different media types (**Figure 9a**). Cells cultured in the HPv7 medium containing double the N content had the greatest stationary cell density; however, the spent medium was acidified to pH 3. One out of three replicate cultures appeared lighter in color, like cells grown in TP +2% (w/v) glucose (**Figure 8)**. The Na-HEPES and potassium phosphate buffer systems used in HPv7 were the same as HPv6, however the pH of the HPv6 medium was maintained at pH 6 (**Figure 8**). HPv1 and HPv4 media were more acidic than HPv6 indicating that the increase in the potassium phosphate and decrease in Na-HEPES buffers were more effective than 20 mM of Na-HEPES and 1.13 mM of total P (**Figure 8**).

**Figure 8.**
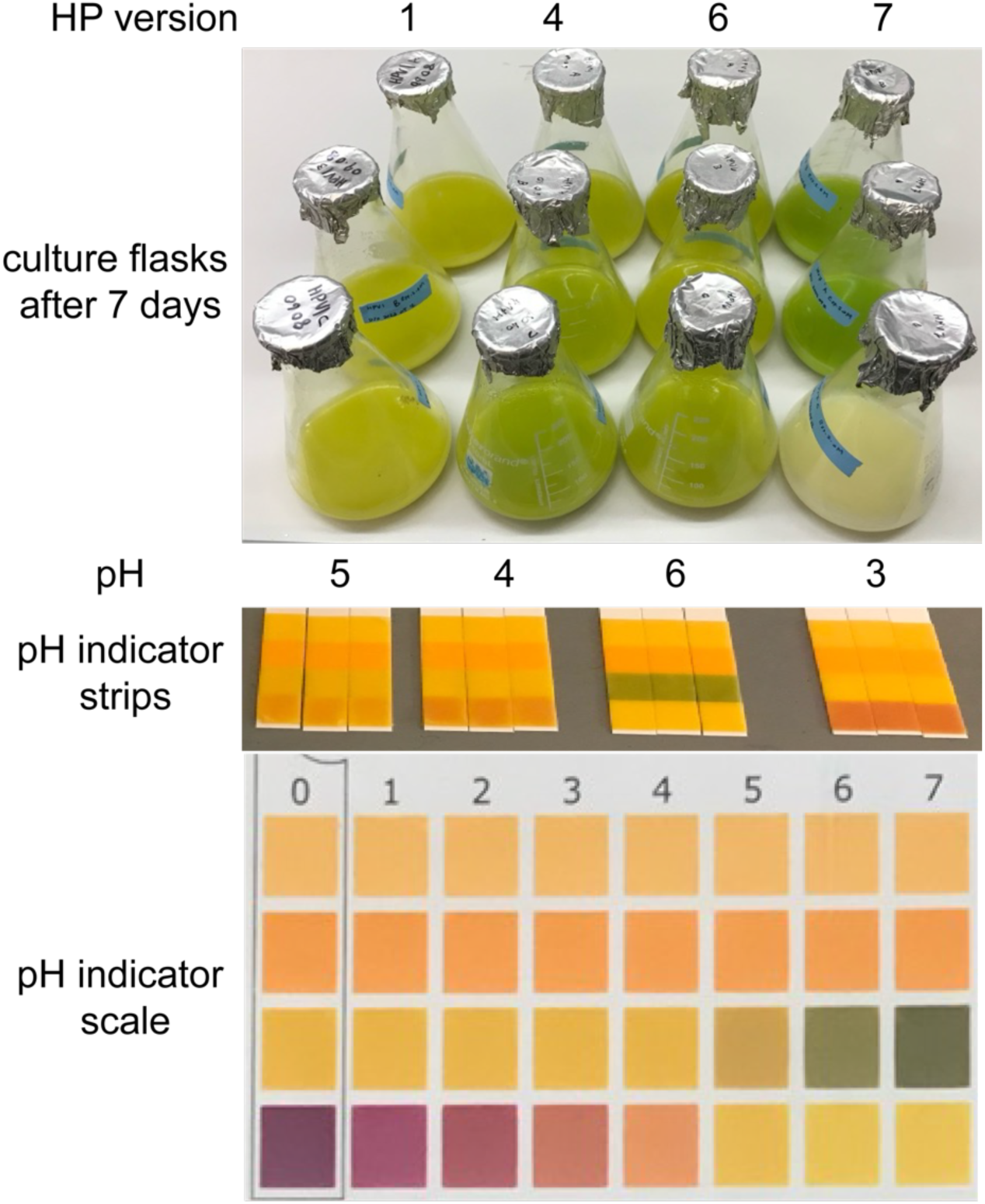
UTEX 250-A culture flasks after 7 days of mixotrophic growth in the specified medium, supplemented with 2% (w/v) glucose and illuminated with 100 µmol photons m^−2^ s^−1^. Three replicate culture flasks were tested for each medium. pH indicator strips displaying the pH of the respective spent medium after 7 days. Cultures were centrifuged at 21,100 ×*g* for 2 min and 30 µL of the supernatant was applied to each strip. The pH indicator scale is shown at the bottom.

**Figure 9.**
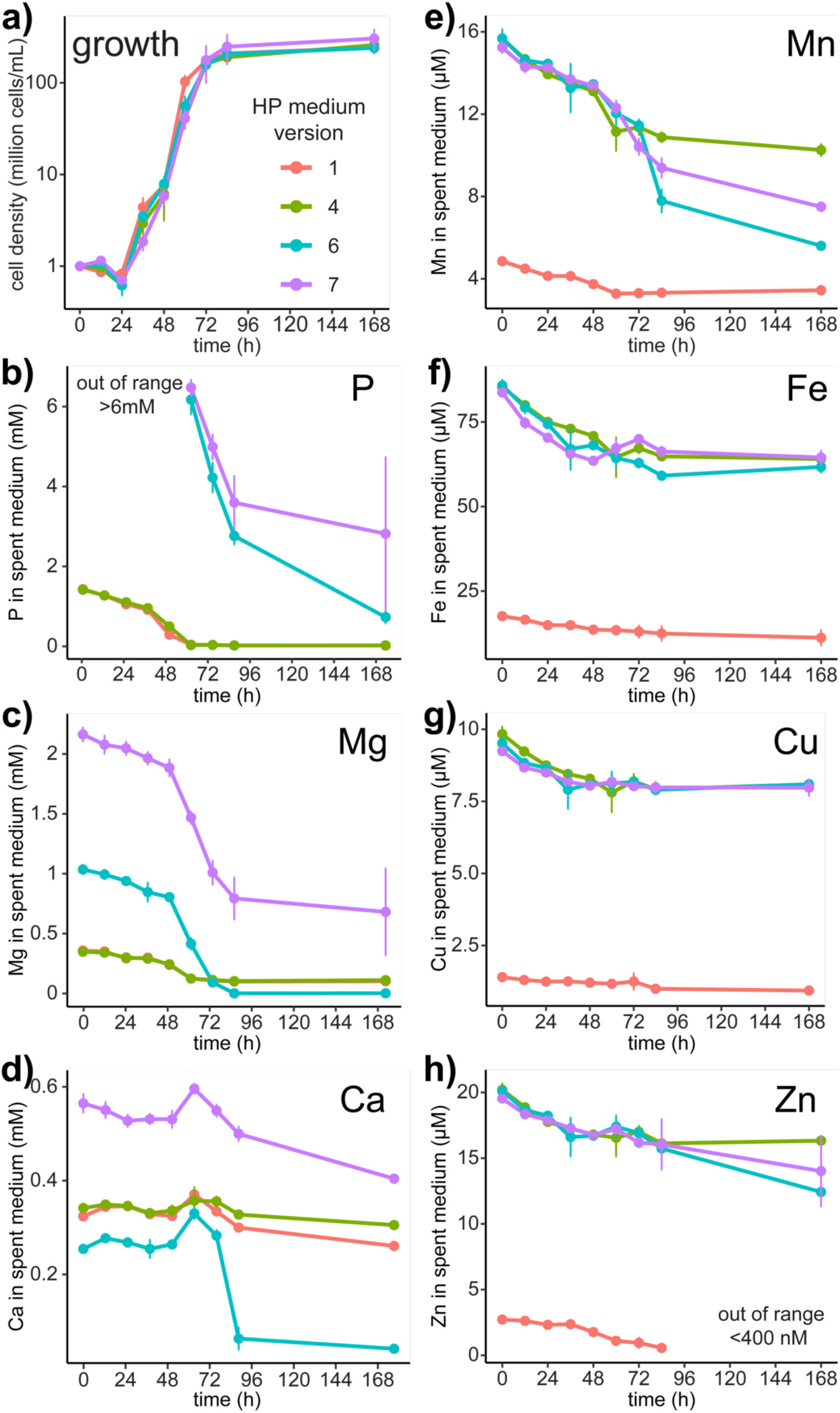
Elemental content of the spent HPv1, HPv4, Hpv6, and HPv7 media during and after UTEX 250-A mixotrophic growth in the light (100 µmol photons m^−2^ s^−1^) with 2% (w/v) glucose supplementation. Error bars represent the standard deviation between three replicate culture flasks.

### Excess phosphorous induces sequential magnesium and calcium depletion in the HPv6 spent medium

Analysis of the spent medium revealed again that P levels in HPv1 and HPv4 were depleted to an average of 35 µM at 60 h, and 22 µM at seven days. However, when P content was increased in HPv6 to prevent complete depletion, sharp declines in Mg, Mn, and Ca levels were observed after 84 h (**Figure 9c,e,d**). A comparison of Mg concentrations between HPv6, HPv1, and HPv4 at 84 h suggests that Mg assimilation was suppressed when P was insufficient in HPv1 and HPv4 but was permitted when excess P was provided in HPv6 (**Figure 9c**). In most cells, up to 50% of cytosolic Mg^2+^ is bound to ATP and the Mg:ATP salt serves as the substrate for most ATP-dependent enzymatic reactions, rather than free ATP anions (Bruna et al., 2021; Maguire & Cowan, 2002; Storer & Cornish-Bowden, 1976). With the increased P content in HPv6, Mg levels in the spent medium plummeted to 2 µM, 50-fold lower than the 100 µM Mg measured in HPv1 and HPv4 at seven days and 500-fold lower than the initial Mg content measured in HPv6, speaking to co-uptake of Mg ions with phosphate (**Figure 9c**). Polyphosphates compartmentalized in acidocalcisomes may chelate Mg (Momeni & Filiaggi, 2014) and act as an intracellular ion filter to extract divalent and trivalent metal ions from intracellular vesicles that originate from fluid phase endocytosis (Klompmaker et al., 2017). Notably, Ca was rapidly assimilated, only after Mg was depleted in HPv6 at 84 h, a response also observed in Mg deficient yeast (Wiesenberger et al., 2007). The Mn level in the spent medium also decreased in HPv6, consistent with co-transport of Mn ions with phosphate (Jensen et al., 2003). In Chlamydomonas, Mn co-accumulates with phosphorous, eventually colocalizing with Ca and polyphosphates in the acidocalcisomes of Chlamydomonas, although Mn eventually forms mononuclear complexes with inorganic P_i_ and phytate (Tsednee et al., 2019).

In contrast to HPv6, the elevated Mg concentration (2.4 mM) in HPv7 remained sufficient through 168 h. Consequently, the Ca and Mn concentrations in HPv7 were maintained relative to the decreases in Ca and Mn concentrations observed in HPv6 at 84 h. The delayed assimilation of Ca and Mn in cells cultivated in HPv6 suggests that their uptake may be mediated by promiscuous divalent cation transporters with higher affinity for Mg, transporting Mn and Ca into the cell only after Mg was depleted.

### Ca and Mn overaccumulation is ameliorated with additional Mg

We then formulated and tested three more versions of the medium simultaneously: HPv8, HPv9, and HPv10 (**Table 3**). Given the potential for excess Fe to catalyze Fenton reactions leading to photo-oxidative stress (Dixon et al., 2012; Halliwell & Gutteridge, 1986; Long & Merchant, 2008), we chose to reduce the Fe content from 100 µM to 20 µM, reverting to the original concentration of Fe in TAP and HPv1. Notably, the Fe content in ApM1 (4 µM) is lower than that of TAP and HPv1 (20 µM). In HPv1, 11 µM of 18 µM Fe measured at the beginning of the experiment remained in the spent medium after 7 d. On average, 60 µM of the 85 µM Fe initially measured in HPv4, HPv6, and HPv7 remained after 7 d. The differences between the three new versions were minimal, with differences in only one component between each successive version, allowing for the systematic dissection of each element’s contribution. Since HPv6 demonstrated adequate buffering capacity, HPv8 was formulated almost identically, with the only modification being a reduction in Fe content (**Tables 3 and2).** HPv9 retained the composition of HPv8 but included an increased MgSO_4_ concentration (2.4 mM) to match HPv7, since the Mg level in the stationary phase HPv7 spent medium did not decrease to HPv1, HPv4, and HPv6 levels. HPv10 remained unchanged from HPv9, apart from the complete omission of Na-HEPES. To minimize the use of unnecessary HEPES buffer containing S and Na, we wanted to evaluate whether the increased potassium phosphate buffer alone was able to maintain the pH of the medium. We previously attempted to omit Na-HEPES completely in HPv3 and HPv5 but were unable to test cell growth using these media due to precipitation (**Figure 7**).

**Table 3.** Composition of HPv8–HPv10. Differences from the preceding versions are highlighted in yellow.

|  | species | TAP | HPv6 | HPv8 | HPv9 | HPv10 |
| --- | --- | --- | --- | --- | --- | --- |
| (mM) | NH<sub>4</sub>Cl | 7.5 | 7.5 | 7.5 | 7.5 | 7.5 |
|  | CaCl<sub>2</sub> | 0.3 | 0.3 | 0.3 | 0.3 | 0.3 |
|  | MgSO<sub>4</sub> | 0.4 | 1.2 | 1.2 | 2.4 | 2.4 |
|  | HEPES | 0 | 10 | 10 | 10 | 0 |
|  | K<sub>2</sub>HPO<sub>4</sub> | 0.7 | 6.8 | 6.8 | 6.8 | 6.8 |
|  | KH<sub>2</sub>PO<sub>4</sub> | 0.5 | 4.5 | 4.5 | 4.5 | 4.5 |
| (μM) | Zn | 2.5 | 20 | 20 | 20 | 20 |
|  | Mn | 6 | 18 | 18 | 18 | 18 |
|  | Fe | 20 | 100 | 20 | 20 | 20 |
|  | Cu | 2 | 15 | 15 | 15 | 15 |
| (nM) | Mo | 28.5 | 28.5 | 28.5 | 28.5 | 28.5 |
|  | Se | 1 | 1 | 1 | 1 | 1 |

We cultivated UTEX 250-A in HPv1, HP8, HPv9, and HPv10 and monitored growth, every 12 h for 120 h (**Figure 11a**). We took elemental measurements of the spent media at three timepoints; before the experiment, 72 h post inoculation, and once more seven days post inoculation. The growth of UTEX 250-A in HPv8–HPv10 media was not enhanced relative to cells simultaneously cultivated in HPv1 (**Figure 11a**). The reduction of Fe from 100 µM to 20 µM had no obvious effect on the assimilation of other nutrients. The Fe content of the spent HPv9 medium was the lowest at 7 d compared to all other HP media compositions (**Figure 11f**). Fe was measured at 14 µM pre-inoculation and at 4 µM 7 d post-inoculation, equivalent to the Fe concentration supplied in fresh ApM1 medium. As seen for growth in HPv6, the spent HPv8 medium also had a substantial decrease in Mg and Ca content at day 7 (**Figure 11d**). The pH of the spent media was measured after 14 days and revealed that the complete omission of Na-HEPES in HPv10 caused the pH to decline to 3 (**Figure 10**). The pH of HPv8 and HPv9, containing 10 mM of HEPES (titrated with NaOH) and 11.3 mM of total P, were maintained at pH 6. The Cu content of the spent HPv9 medium was also the lowest at 7 d, in comparison to spent HPv8 and HPv10 media. In HPv9, Cu was measured at 8 µM at 0 h and at 7 µM in the spent medium 7 d post inoculation. The only difference between HPv8 and HPv9 was the increase of Mg from 1.2 mM to 2.4 mM, which attenuated the cellular accumulation of excess Mn and Ca (**Figure 11de**). At day 7, the spent HPv9 medium contained 2 µM more Mn than HPv8 (**Figure 11e**). Similarly, the Mn levels in the Mg depleted spent HPv6 medium were lower than in spent HPv4 and HPv7 media (**Figure 9e**). However, it is unclear why Mn levels are maintained in the spent HPv10 medium relative to HPv8 and HPv9 spent media.

**Figure 10.**
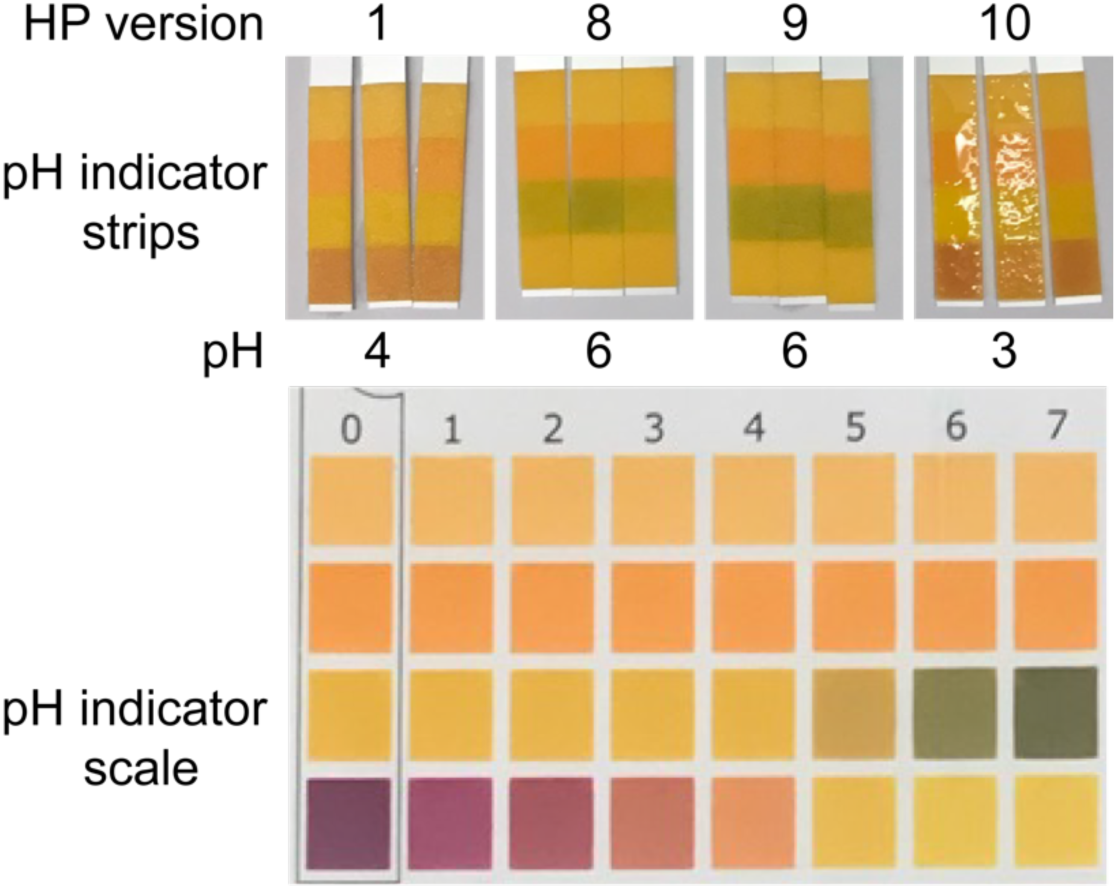
pH measurement of the spent medium 14 days post inoculation. 50 µL from each culture was centrifuged at 21,100 ×*g* for 2 min and 30 µL of the supernatant was applied to each pH indicator strip. A pH indicator strip from each of the three replicate culture flasks for each medium type is shown.

**Figure 11.**
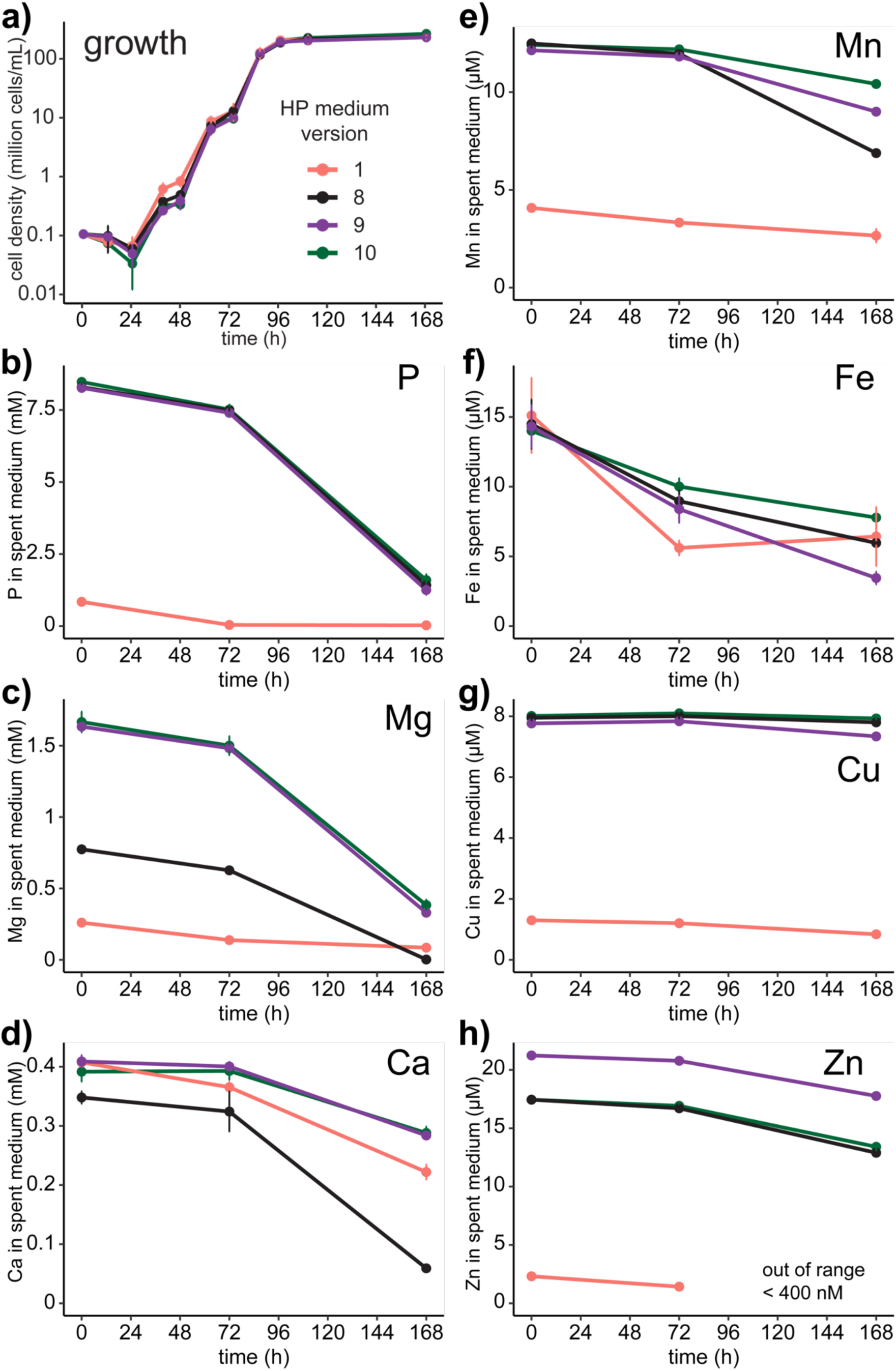
Growth and nutrient consumption of *Auxenochlorella* (UTEX 250-A) in HPv1, HPv8, HPv9, and HPv10 supplemented with 2% (w/v) glucose. Error bars represent the standard deviation from the average of three replicate culture flasks for each medium. **a)** Growth curve of UTEX 250-A after seven days in illuminated incubators (100 µmol photons m^−2^ s^−1^), at 28 °C, and agitated at 200 RPM. Cells were counted every 12 h using a hemocytometer. **b–h)** Concentration of elements present in each medium measured before the experiment, 72 hours post inoculation, and 7 days post inoculation.

**Figure 12.**
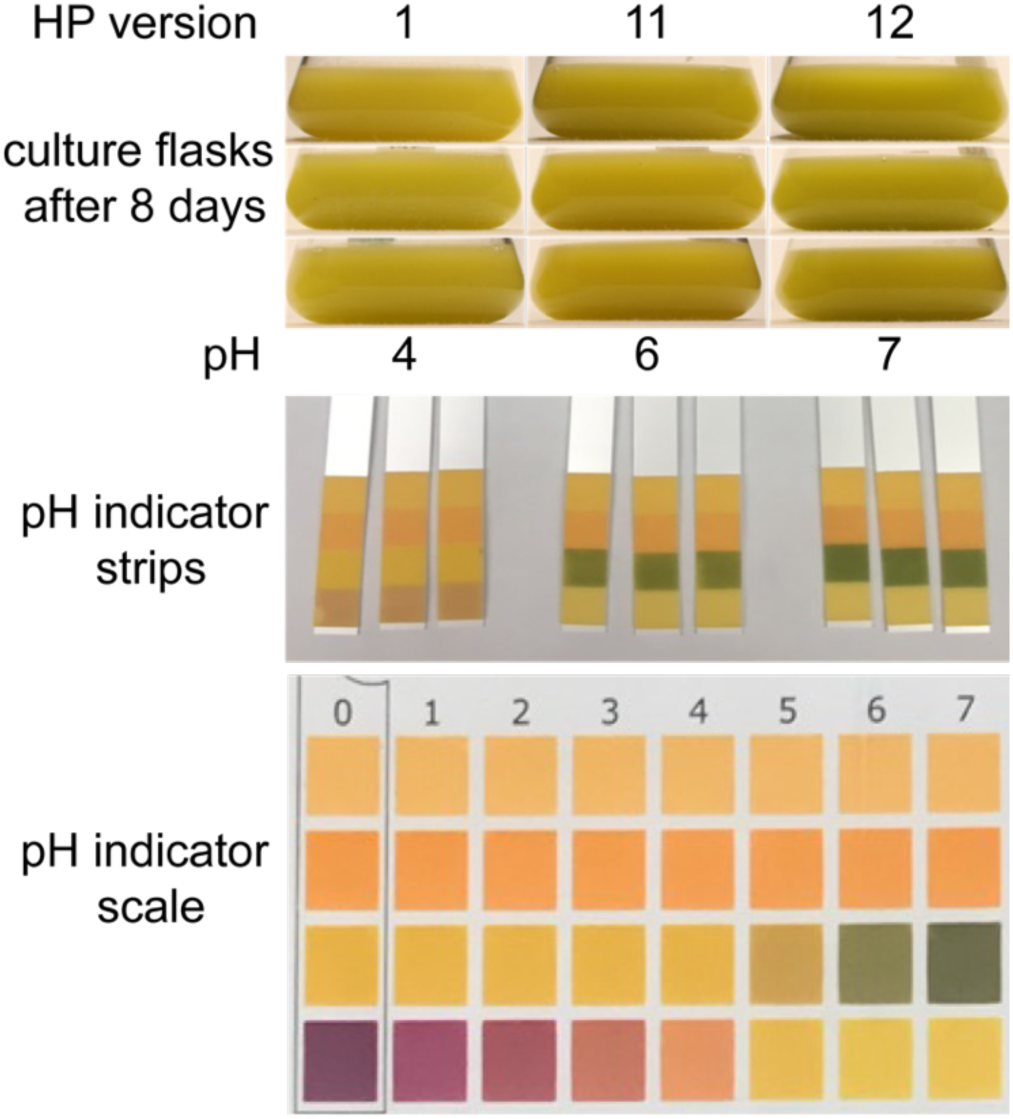
Culture flasks and pH of the spent HPv1, HPv11, and HPv12 media 8 d post inoculation. 50 µL from each culture was centrifuged at 21,100 ×*g* for 2 min and 30 µL of the supernatant was applied to each pH indicator strip. A pH indicator strip from each of the three replicate flasks for each medium type is shown.

### The HPv11 medium is replete

In the third round of optimizations, we used HPv9 as a chassis for the formulation of HPv11 and HPv12. Given the consumption of only 1 µM Cu by cells cultivated in HPv9, we chose to lower the Cu concentration of HPv11 to 2 µM, equivalent to the Cu concentration in TAP. We also used the same buffer system as HPv6–HPv9 for HPv11. The Zn contents of HPv8–HPv10 were meant to be provided at equal concentrations, however HPv9 was closer to the target of 20 µM than were HPv8 and HPv10. Nonetheless the amounts of Zn assimilated were very similar (**Figure 11h**). On average, cells cultured in HPv8–HPv10 removed 4 µM Zn ions from the medium. We chose to lower the Zn content from 20 µM to 10 µM because it was harder to deprive cells of Zn in preliminary Zn deficiency experiments when we maintained them in 20 µM.

To formulate HPv12, we calculated the difference of HPv9’s elemental composition before and after 7 d of growth and then multiplied it by 3 (**Equation 2**). We provided approximately 3-fold excess of each element by raising the concentration of MgSO_4_ to 4 mM, P to 22.6 mM, Fe to 30 µM Fe, and Cu to 2.5 µM in HPv12 (**Table 4**). Following the same strategy employed in the previous experiments, we cultivated the algae in HPv1, HPv11, and HPv12 media simultaneously and measured their cell densities approximately every 12 h for 96 h and at 8d post-inoculation. The spent media were measured every 12 h for 96 h except for a timepoint at 72 h.

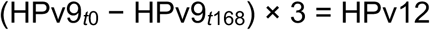

**Equation 2.** HPv12 formulation using the cellular uptake of elements from the HPv9 medium.

**Table 4.** Composition of HPv1, HPv11, and HPv12. Modifications to the previous medium are highlighted in each new medium.

|  | species | HPv1 | HPv11 | HPv12 |
| --- | --- | --- | --- | --- |
| (mM) | NH<sub>4</sub>Cl | 7.5 | 7.5 | 7.5 |
|  | CaCl<sub>2</sub> | 0.3 | 0.3 | 0.3 |
|  | MgSO<sub>4</sub> | 0.4 | 2.4 | 4 |
|  | HEPES | 20 | 10 | 10 |
|  | K<sub>2</sub>HPO<sub>4</sub> | 0.7 | 6.8 | 13.6 |
|  | KH<sub>2</sub>PO<sub>4</sub> | 0.5 | 4.5 | 9 |
| (μM) | Zn | 2.5 | 10 | 10 |
|  | Mn | 6 | 12 | 12 |
|  | Fe | 20 | 20 | 30 |
|  | Cu | 2 | 2 | 2.5 |
| (nM) | Mo | 28.5 | 28.5 | 28.5 |
|  | Se | 1 | 1 | 1 |

The addition of excess Mg, S, K, P, Fe, and Cu in HPv12 did not enhance the growth of UTEX 250-A (**13a**), and the net uptake of most elements remained similar to HPv11 (**Figure 14**). The assimilation of P, Mn, Cu, and Zn did not differ significantly between HPv11 and HPv12. Although the differences of net assimilation of Mg, Ca, and Fe were significant between HPv11 and HPv12, Mg, Ca, and Fe remained at sufficient levels in HPv11 after eight days. The average Mg concentration of the spent HPv11 spent medium after eight days was measured at 0.2 mM (**Figure 13c**), half the amount supplied in a replete TAP and HPv1 medium (0.4 mM). Although the assimilation of Ca was greater in cells grown in HPv11 compared to HPv12, it was comparable to cells grown in HPv1, and substantially lower than the net assimilation observed in cells grown in HPv6 and HPv8 (**Figure 15**). The average Ca content of the spent HPv11 medium after 8 days was 150 µM (**Figure 13d**), more than double the amount remaining in spent HPv8 (60 µM Ca, **Figure 11d**) and triple the amount in spent HPv6 (40 µM Ca, **Figure 9d**) measured seven days post inoculation. The Fe concentration of spent HPv11 eight days post inoculation was measured at 4 µM (**Figure 13f**), equal to the Fe concentration supplied in a replete ApM1 medium. Taken together, these observations suggest that the additional supplementation in HPv12 did not confer any physiological advantage over HPv11, and that HPv11 was already replete, as all elements tested remained sufficiently available in spent media even after eight days.

**Figure 13.**
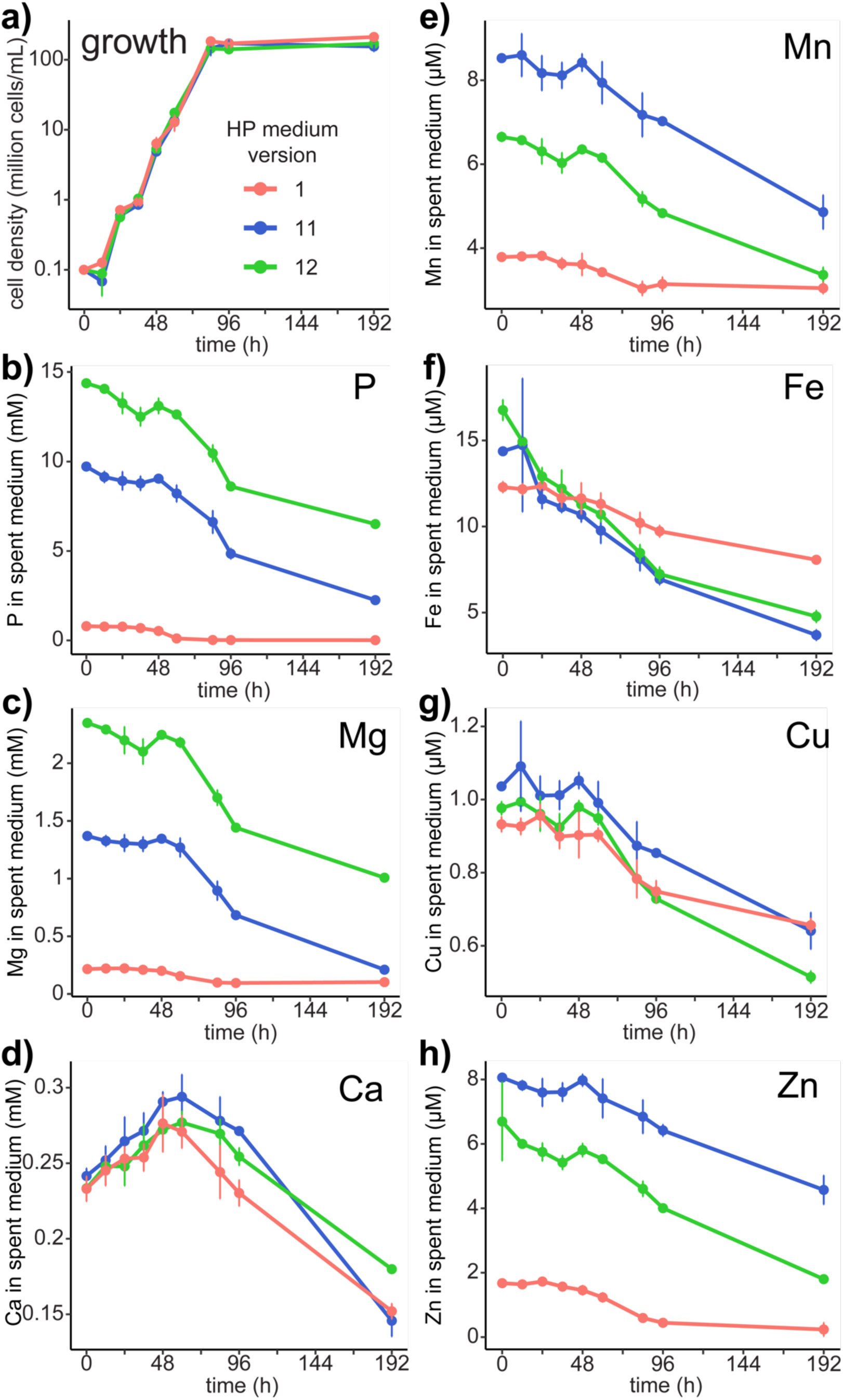
Growth and nutrient consumption of *Auxenochlorella* (UTEX 250-A) in HPv1, HPv11, and HPv12 supplemented with 2% (w/v) glucose. Error bars indicate standard deviations from the average of three replicate flasks cultivated in each medium. **a)** Growth curve of UTEX 250-A after eight days in illuminated incubators (100 µmol photons m^−2^ s^−1^), at 28 °C, and agitated at 200 RPM. Cells were counted every 12 h using a hemocytometer. **b–h)** Concentration of elements present in each spent medium measured every 12 h for 96 h except for at 72 h and again after eight days.

**Figure 14.**
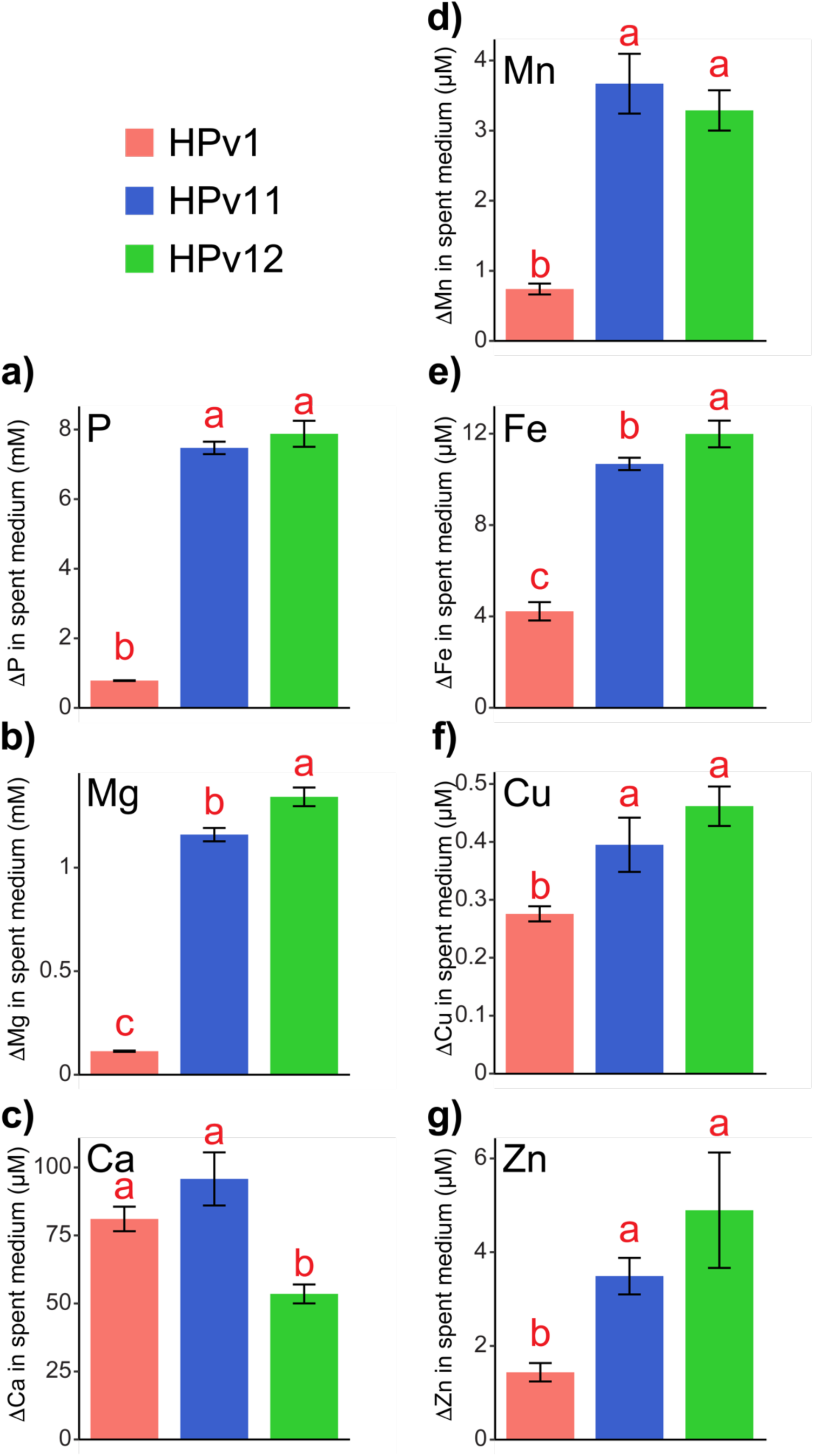
Average net uptake of elements by UTEX 250-A cells over eight days (192 h). Three replicate flasks were cultivated for each medium type. Uptake was calculated by subtracting the concentration of elements in the stationary phase (192 h) spent medium from the initial medium measured at 0 h. Differences were calculated for each individual replicate and the averaged difference is plotted. Error bars represent the standard deviation between the differences calculated for each specific replicate. Elemental uptake was compared between conditions using a one-way ANOVA test followed by a Tukey’s Honest Significant Difference (HSD) post-hoc test (a = 0.05). Red letters above the bars indicate statistically significant differences.

**Figure 15.**
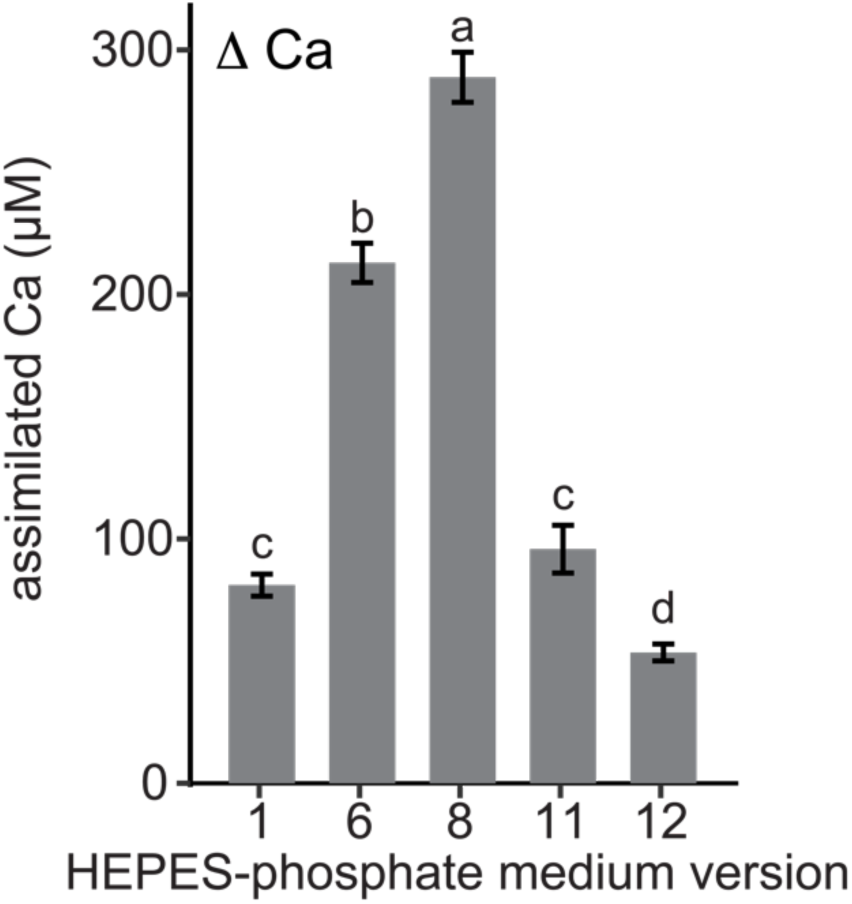
Comparison of Ca assimilation by UTEX 250-A cells cultivated in different versions of the HP medium. Three replicate flasks were cultivated for each version. Assimilation was calculated for each version by subtracting the Ca content in stationary phase spent medium from the initial medium. Stationary phase spent media of HPv1, 11, and 12 were collected at 192 h post inoculation while HPv6 and HPv8 spent media were collected at 168 h. The differences for each individual replicate were calculated and the averaged differences are plotted. Error bars represent the standard deviation between differences calculated for each replicate. Statistically significant differences are denoted using letters above the bars and were determined using one-way ANOVA then a Tukey’s Honest Significant Difference (HSD) post-hoc test (a = 0.05).

### The systematic modulation of specific elements in the HPv11 medium uncovers a link between P assimilation and the mixotrophic to autotrophic transition

The elemental compositions of HPv1, HPv4, HPv6, and HPv8 suggested a deficiency of one or more nutrients, however none of the other versions enhanced growth. The exception was HPv7, which stimulated a higher stationary cell density, but suffered from acidification due to the increased NH_4_ content. This raised the question of whether the medium lacked a crucial element or if certain elements were provided at suboptimal ratios relative to others, causing toxicity. To determine whether HPv11 was truly a replete medium, we cultivated UTEX 250-A in 20 different versions of HPv11, each with a single targeted modification. We included a version with a five-fold increase of C (10% (w/v) glucose) and a separate version with a five-fold increase of NH_4_ to assess any potential improvements to growth. The following data should be interpreted with caution as only a single replicate for each condition was experimentally tested. The goal of this approach was to obtain a broad survey of the growth phenotypes and characteristic elemental profiles associated with either a deficiency or an over supplementation of a given nutrient.

The growth and elemental content were monitored every 24 h for 96 h and a final timepoint was taken at 7 days post inoculation to assess the stability of cells at stationary phase. Distinct visual phenotypes associated with some deficiencies and toxicities were observed after 48 h (**Figure 16**) The “replete” HPv11 culture is green at 48h and becomes yellow as it transitions into stationary phase after 72 h. When all glucose was presumably consumed, the culture regreened as it transitioned to photoautotrophic growth. When no glucose was provided, the culture did not turn yellow and remained green. When 5-fold glucose was provided, the culture did not regreen after 7 days. A sudden decrease in the P content of the replete medium relative to the P content of C limited and excess (5×C) media at 96 h indicates that the trophic switch is responsible for the increased P demands of the cell, rather than photosynthesis itself (**Figure 17**). The same pattern was observed in all cultures that greened after 7 days (replete, 5×S, 5×Ca, 5×Fe, 5×Cu, −Zn, 5×Zn, 5×Mn, 5×Mo, and 5×Se) and was not observed in cultures that were unable to make the trophic transition (**Figure 18**) as determined only visually by the color of the culture at 168 h (**Figure 16**). We observed no other indications of nutrient insufficiency besides C limitation in HPv11.

**Figure 16.**
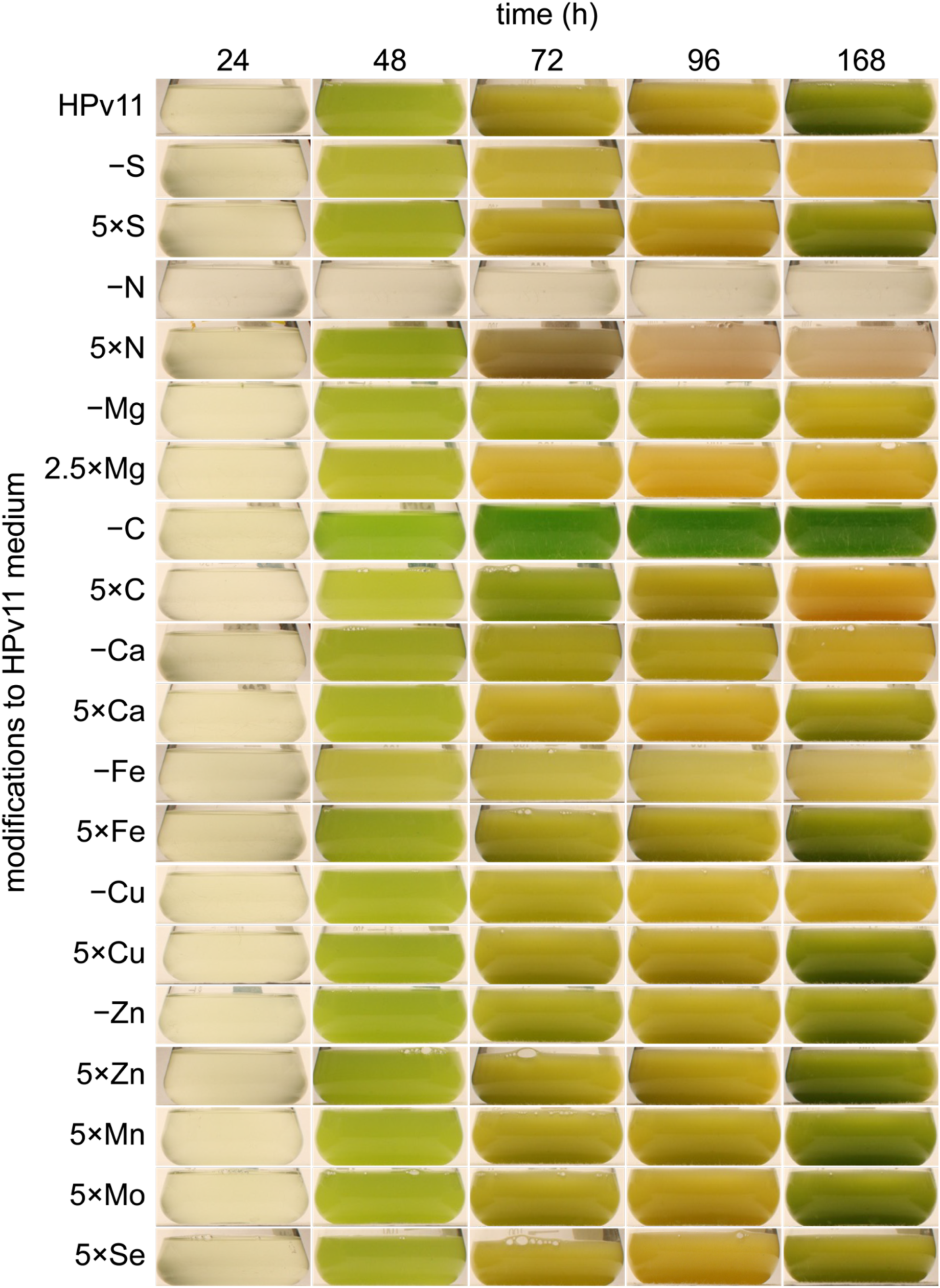
UTEX 250-A culture flasks throughout growth in HPv11 medium supplemented with 2% (w/v) glucose (except for −C and 5×C). A single modification to the medium is indicated to the left.

**Figure. 17.**
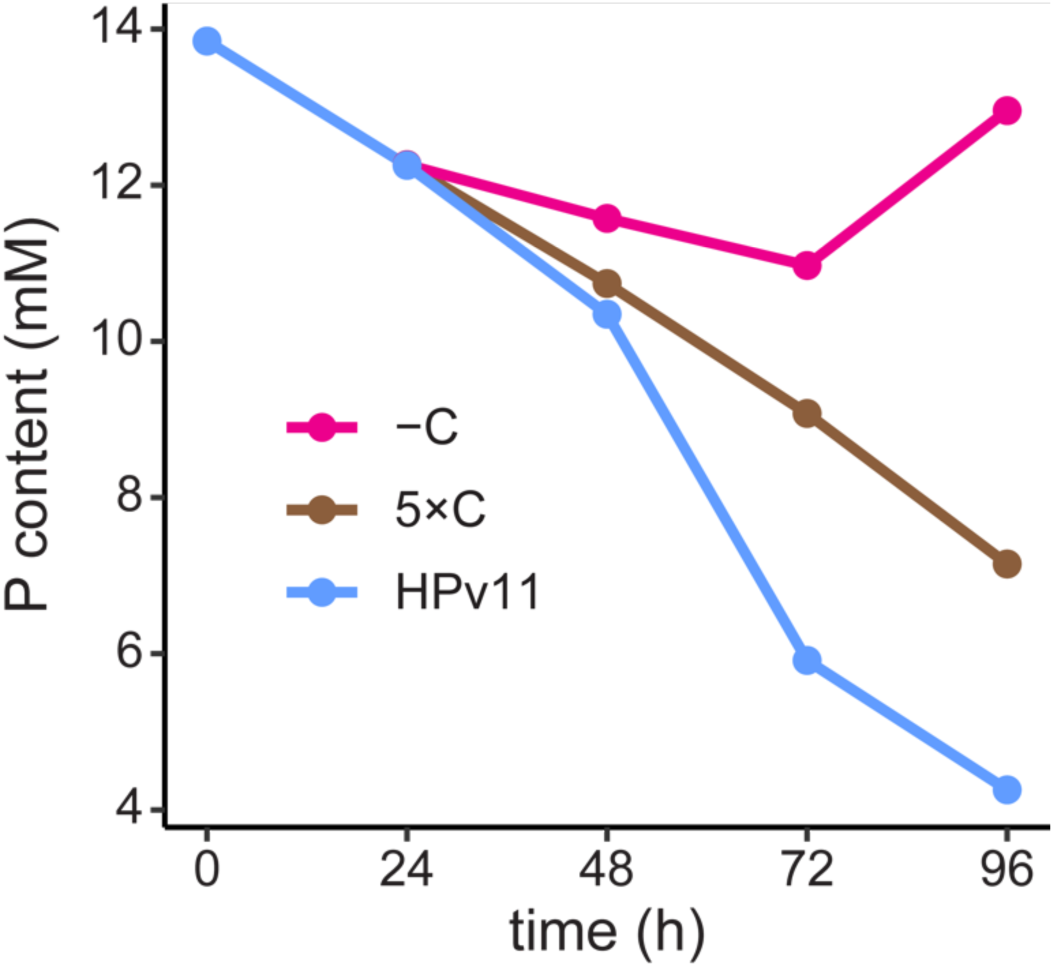
Phosphorous content of spent HPv11 medium with varying levels of glucose during growth of UTEX 250-A in illuminated incubators (100 µmol photons m^−2^ s^−1^), at 28 °C, and agitated at 200 RPM. The normal HPv11 (blue line) was supplemented with 2% (w/v) glucose. 5×C represents a 5-fold increase in glucose to 10% (w/v) glucose, and −C cultures were not supplemented with glucose. One replicate flask was measured for each condition.

**Figure 18.**
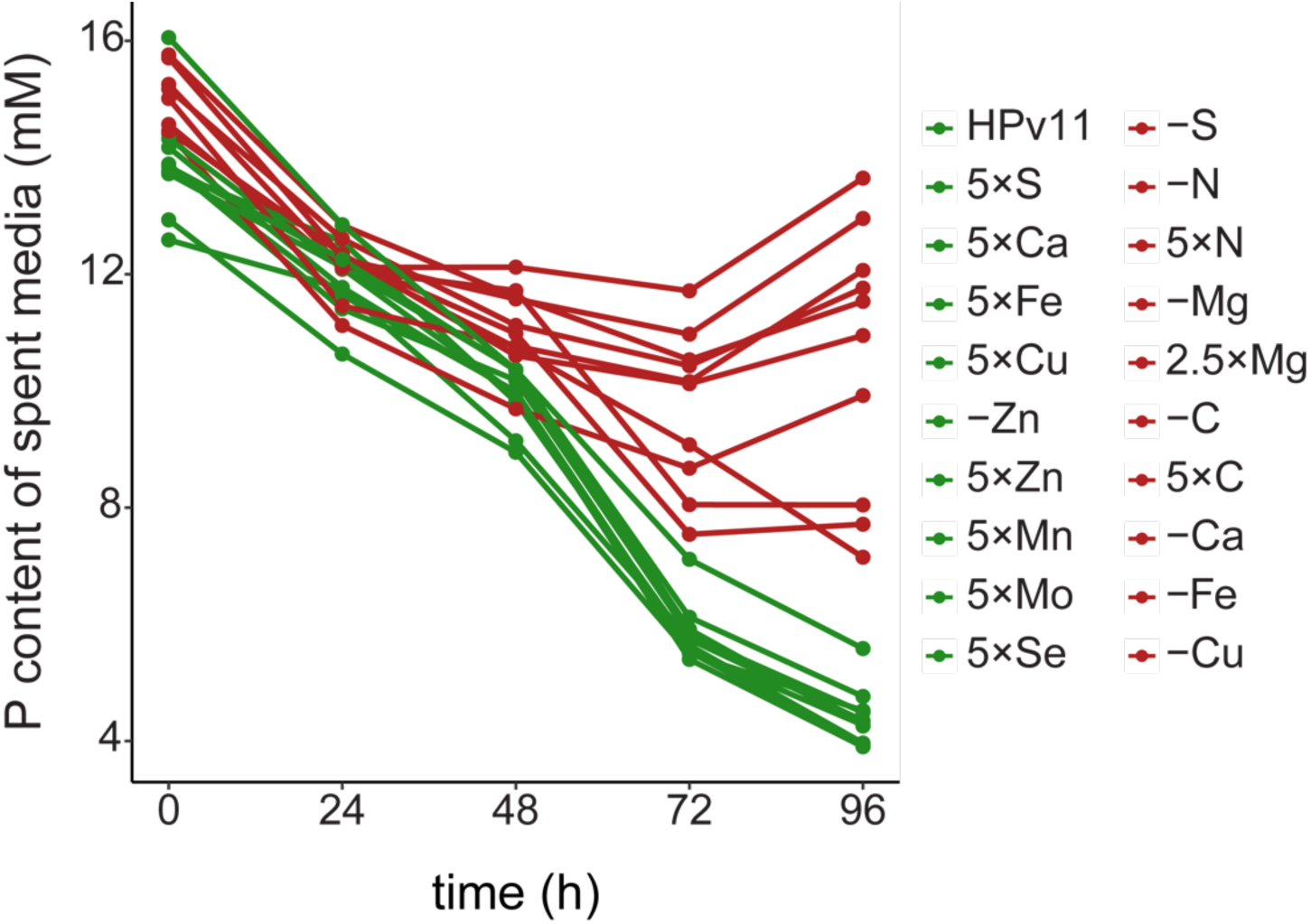
Phosphorous content of all spent media types tested during growth of UTEX 250-A. Green lines represent the P content of media types where the cultures transitioned to photoautotrophic growth after 168 h determined by the color of the cultures (Figure 16). Red lines represent media types that failed to transition to photoautotrophic growth (Figure 16). Phosphorous content from one replicate flask per nutrient condition was measured.

## Conclusions

We outlined the formulation and optimization of a replete UTEX 250-A culture medium, adapting the widely used TAP medium for mixotrophic growth in 2% glucose. We observed acidification of the HSM and TP spent media when UTEX 250-A cells were supplemented with 2% glucose. To restore pH balance we replaced TRIS with a more suitable buffer, Na-HEPES. Next, we monitored the elemental content of the spent medium throughout UTEX 250-A’s growth and determined that cells were experiencing P, Mg, Mn, and Zn deficiencies. We raised the P content of the medium and took advantage of the buffering capacity provided by the monobasic and dibasic mixture of potassium phosphate to lower the amount of Na-HEPES that had to be provided for buffer capacity. The increase in P content of the medium resulted in a greater capacity for cellular Mg assimilation. When the Mg content of the medium was depleted, cells began to assimilate Ca. Mn levels in the spent medium also decreased simultaneously. The provision of these nutrients in subsequent versions of the medium failed to significantly enhance growth. The majority of P, Mg, Ca, and Mn assimilation occurred after cells had reached stationary phase, indicating that these excess nutrients were not necessary for growth, but rather for stabilization at the stationary phase. A survey of controlled deficiency and toxicity induced phenotypes conducted in the HPv11 medium only found that the depletion of P content in the replete spent medium was attributed to a trophic switch necessitated by C limitation. The HPv11 medium is compatible with ICP-MS instrumentation and is suitable for the systems level analysis of nutrient deficiencies in UTEX 250-A. A detailed protocol describing the preparation of the final medium for trace metal studies is publicly available (Camacho, Perrino, et al., 2024c). Transcriptomes generated from some of the nutrient deficiencies tested in this study were used for the gene model predictions and structural annotation of the UTEX 250-A genome (Craig et al., 2025). The annotation of genes exclusively expressed during these conditions will aid gene model dependent comparative transcriptomics and proteomics analysis of UTEX 250.

## Acknowledgments

We thank Liam Sheedy and Marco Duenas for feedback on the manuscript. We thank Dr. Radhika Mehta for guidance and training on TEM sample preparation and imaging. We are grateful to Dr. Daniel Jorgens and Reena Zalpuri at the University of California Berkeley Electron Microscope Laboratory for their advice and assistance in electron microscopy sample preparation and data collection. This work was supported by two grants from the National Institutes of Health (NIH): grant no. GM 042143 (for work on Cu) and grant no. 5T32GM007232-44 (training grant for Dimitrios J. Camacho). It was also supported by two grants from the US Department of Energy (DOE), Office of Biological and Environmental Research (BER): grant no. DE-SC0020627 (for work on Fe) and grant no. DE-SC0023027 (for work on Auxenochlorella (UTEX 250-A). Dimitrios J. Camacho was also supported by the University of California, Berkeley, Chancellor’s Fellowship.

